# Equilibrative Nucleoside Transporter 3 is an IFN-stimulated Gene that Facilitates Viral Genome Release

**DOI:** 10.1101/2022.04.27.489746

**Authors:** Yu-Ting Hsieh, Tsung-Lin Tsai, Chia-Chun Tu, Shen-Yan Huang, Jian-Wen Heng, Pei-Yuan Tsai, Tai-Ling Chao, Ya-Min Tsai, Pei-Ching Chang, Chien-Kuo Lee, Guann-Yi Yu, Sui-Yuan Chang, Ivan L. Dzhagalov, Chia-Lin Hsu

## Abstract

An increasing body of evidence emphasizes the role of metabolic reprogramming in immune cells to fight off infections. However, little is known about the regulation of metabolite transporters that facilitate and support metabolic demands. In this study, we found that equilibrative nucleoside transporter 3 (ENT3) expression is part of the innate immune response, and is rapidly upregulated upon bacterial and viral infection. The transcription of ENT3 is directly under the regulation of IFN-induced signaling, positioning this metabolite transporter as an Interferon-stimulated gene (ISG). Moreover, we unveil that several viruses, including SARS-CoV2, require ENT3 to facilitate their entry into the cytoplasm. The removal or suppression of ENT3 expression is sufficient to significantly decrease viral replication in vitro and in vivo.

## Introduction

Upon infection, the immune system must recognize the threat to initiate necessary responses, remove the insults, and regain homeostasis. The fast-acting innate immunity is responsible for the initial recognition and early responses and is vital to recruit antigen-specific adaptive immunity. The longstanding co-evolution between the immune system and pathogens has driven innate immune cells to generate various arsenals fighting off the pathogens; on the other hand, the pathogens also continuously develop numerous ways to hide from or interfere with the immune responses. Once the pathogen invasion is established, responding cells shift from quiescence to activation and reprogram their metabolism to support the rapid cell division and production of effector molecules. When the threat is eliminated, the immune system readjusts itself to another metabolic setting allowing the recycling and redistributing of the surplus biomaterials generated from these retired effector cells. Eventually, the metabolic program switches back toward the dormant state and reclaims homeostasis. Multiple steps and various molecules are essential to maintain such versatile yet coordinated programs. Among them, the metabolite transporters play a vital part in facilitating the influx or efflux of nutrients.

The importance of metabolite transporters in mediating cell survival and function has recently received more attention, especially in the immunology and cancer biology fields. It is perhaps not surprising that the vital roles of metabolite transporters were first revealed in vigorously proliferating cells such as activated immune cells and tumor cells. It has become clear that the availability of nutrients is key to immune cell function, growth, and proliferation. Taking T cells, for example, upon activation, various metabolic pathways are activated to meet the biosynthetic needs during the effector phase. The acquisition of the building blocks relies on the metabolite transporters(Hsu & Dzhagalov, 2019). The effector T cells express glucose transporter 1 (Glut1) to obtain glucose, various amino acid transporters for amino acid uptake(Wang & Zou, 2020), and G protein-coupled receptors (GPCRs), CD36, and the fatty acid transport protein (FATP) to attain fatty acids(Howie *et al*, 2018). Together, these transporter proteins support the metabolic demands during the activation and differentiation stage of effector T cells. The expression of metabolite transporters is also a prerequisite for the differentiated myeloid cells or B cells to perform their functions. Glut1 is indispensable for the inflammatory response of macrophages(Freemerman *et al*, 2014); Zinc transporter participates in the phagocytosis and cytokine production of dendritic cells (DCs) and macrophages(Hall *et al*, 2021) as well as the B cell development(Anzilotti *et al*, 2019). These observations demonstrate the importance of metabolite transporter and immune effectors and raise the interesting question of how the sensing of threat is translated into metabolic reprogramming. Immune cells constantly survey and respond to environmental cues, including microbial pattern, pH, oxygen availability, or mechanical stiffness(Chakraborty *et al*, 2021). However, the signaling events that regulate metabolite transporters expression or post-translational modification are still underexplored.

Nucleotides are constantly needed in every living cell for the synthesis of DNA, RNA, enzyme co-factors, or signaling molecules. While how cells acquire glucose, amino acids, or fatty acids is under intense investigation, we know relatively little about how the nucleotides are obtained. There are two ways our cells obtain nucleotides – the salvage and de novo nucleotide synthetic pathways. The de novo synthesis is an energy-intensive multistep process that involves several metabolic pathways. In contrast, the salvage pathway recycles the nucleobases and nucleosides from nucleotides degradation and serves as the significant supply for cellular nucleotides pools. Cells use transporter proteins to translocate nucleobases and nucleosides through membranes to salvage these valuable biological building blocks. These membrane proteins are known as equilibrative nucleoside transporters (ENTs). ENTs are metabolite transporters evolutionary conserved from protozoa to humans(Landfear, 2011), belonging to the solute carrier 29 family (SLC29). Thus far, four ENTs (ENT1 to 4) encoded by slc29a1 to 4 have been identified in mammals(Young *et al*, 2013b). ENT1, 2, 4 are positioned at the plasma membrane, while ENT3 has unique intracellular localization and pH-dependency(Young *et al*, 2013a; Baldwin, 2005). ENT1 facilitates the entry of nucleoside analogs for viral and tumor therapies(Jordheim *et al*, 2013), and its mutation or expression level has been associated with the prognosis of cancer treatment efficacy(Macanas-Pirard *et al*, 2017; Takagaki *et al*, 2004). Inhibition of ENT2 has been proposed as a path to resolve intestinal inflammation(Aherne *et al*, 2018) and neuroinflammation(Wu *et al*, 2020). Mutations in ENT3 are identified in multiple inherited diseases in humans, including H syndrome(Priya *et al*, 2010), Faisalabad histiocytosis(FD)(Elbarbary *et al*, 2012), pigmentary hypertrichosis and non-autoimmune insulin-dependent diabetes mellitus (PHID) syndromes(Cliffe *et al*, 2009), Rosai-Dorfman disease(Morgan *et al*, 2010), arthritis and systemic inflammation(Jaouadi *et al*, 2018). While patients with ENT3 mutations display a broad spectrum of clinical symptoms, immune involvement is a shared feature. These results together argue for the vital roles of ENTs in the immune system.

Using a murine knock-out model, we have shown that ENT3 is crucial for the function and homeostasis of macrophages(Hsu *et al*, 2012), and the survival of T lymphocytes(Wei *et al*, 2018). The absence of ENT3 impacts the lysosome-mediated activities, including the turnover of M-CSF/M-CSFR signaling(Hsu *et al*, 2012), hyper-activated TLR7 responses(Shibata *et al*), as well as mitophagy(Wei *et al*, 2018) and autophagy(Nair *et al*, 2019). Although having a significantly increased number of myeloid cells, ENT3^-/-^ mice are more susceptible to Listeria monocytogenes infection, and their macrophages are less effective in bacteria-killing than WT littermates(Hsu *et al*, 2012). Similarly, the histiocytosis-harboring ENT3 mutant patients have been reported to have recurrence of skin and ear infections(Campeau *et al*, 2012) or rhinoscleroma(Bolze *et al*, 2012). Together, these results imply the strong association between ENT3 and the effective pathogen clearance.

In this study, we show that the expression of ENT3 is responsive to the presence of pathogens as part of the innate immune response. Upon bacterial infection, ENT3 expression is induced by LPS-TLR4 via MyD88-independent, TRIF-dependent pathway. We further identify that the regulation of ENT3 is mediated via the Type I IFN-IFNAR axis and the direct binding of STAT1 to the promoter region of Slc29a3. These results suggest ENT3 is a metabolite transporter that belongs to interferon-stimulated gene (ISG). Moreover, we found that viruses can utilize ENT3 to facilitate their release of genome. The removal or decrease of ENT3 expression is sufficient to impact viral replication and enhance virus-challenged animals’ survival significantly. Our findings together provide novel insights into the pathogen-host interactions and how viruses can induce the expression of host genes beneficial to their own life cycle.

## Results

### The expression level of ENT3 in macrophages is responsive to bacterial insults

To investigate whether ENT3 levels are dynamically controlled upon infection, we first challenged bone marrow-derived macrophages (BMDMs) with live *E. coli* and measured the expression level of *Slc29a3* (Supplemental Fig 1). We observed a clear induction of *Slc29a3* starting at 6 hours post-infection and lasting at least for 24 hours (Fig 1A). Clearance of infection by macrophages can be divided into four steps - sensing the presence of pathogens (e.g., microbial peptides/metabolites), recognizing the insult via ligand-receptor interaction, ingesting the pathogens via phagocytosis, and removing it by degradation. To find out if phagocytosis of the bacteria is a prerequisite for *Slc29a3* induction, we applied the phagocytosis inhibitor Cytochalasin D, together with the *E. coli* to BMDMs. No significant differences were found between Cytochalasin D-treated and the control group (Fig 1B and Supplemental Fig 2), ruling out the internalization of *E. coli* as a necessary step to initiate *Slc29a3* expression. To pinpoint the potential microbial products or pathways involved, we tested the induction efficiency of live, heat-killed, or gentamycin-killed bacteria, or live bacterial metabolites. We found that while none of the tested treatments could elicit the same level of *Slc29a3* expression as live *E. coli*, they all consistently provoked mild *Slc29a3* upregulation (Fig 1C), implying that the induction of *Slc29a3* by live *E. coli* is possibly a combined effect of bacteria and their metabolites. A shared component among different stimulants is LPS. We confirmed that LPS alone was sufficient to upregulate *Slc29a3* in BMDMs (Fig 1D). These results suggest that the sensing of LPS is sufficient to induce the expression of the metabolite transporter, ENT3, in macrophages.

**Figure 1.**
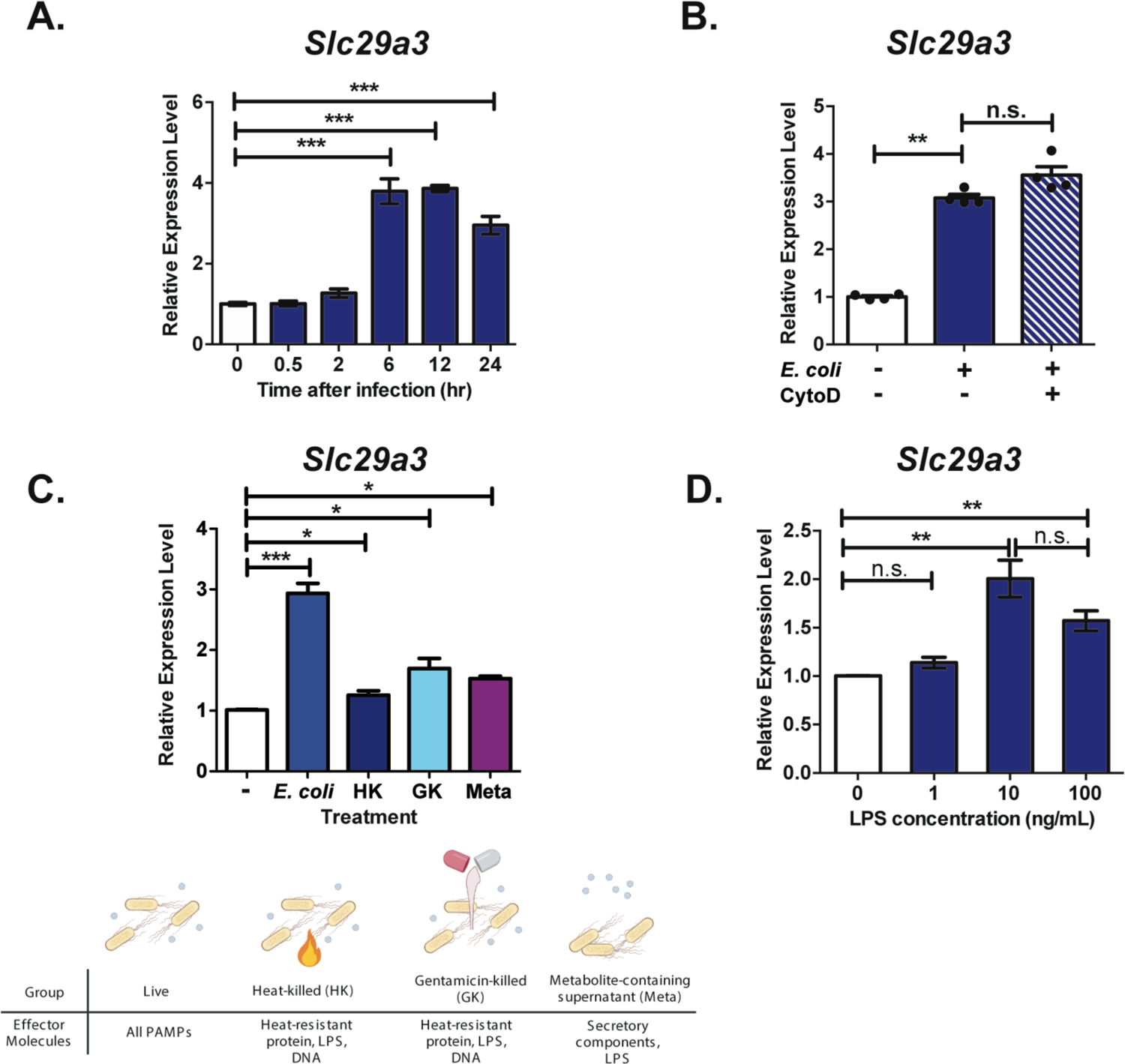
LPS-activated pathways upregulate the expression of ENT3 in BMDMs. **(A)** BMDMs were treated with *E. coli* (MOI=100) for 30 min and further cultured for various length of time as indicated. The dynamic of ENT3 expression was determined by measuring *Slc29a3* transcripts via RT-qPCR. **(B)** Phagocytosis inhibitor, Cytochalasin D (CytoD), pretreated BMDMs were incubated with *E. coli* (MOI=100) and subjected for *Slc29a3* expression analysis at 6 hr post-infection. **(C)** Different bacterial components – live, heat-killed (HK), gentamicin-killed (GK) *E. coli* or metabolite-containing supernatant (Meta), were prepared and applied as stimulants to BMDMs. *Slc29a3* expression was analyzed after 30 min of stimulation and a total of 6 hr incubation. **(D)** BMDMs were treated with different doses of LPS for 30 min, harvested 6 hr post-stimulation and subjected for RT-qPCR analysis. The expression level was calculated relative to *Rpl19* and normalized to untreated group as 1. The results shown are combined from three independent experiments (N=3), unpaired two-tailed Student’s *t*-test was used for statistical analysis, \**p* < 0.05, \*\**p* < 0.01, \*\*\**p* < 0.001, n.s. no significance.

### MyD88-independent pathway regulates the expression of *Slc29a3* in macrophages

As one of the most well-studied pathogen-associated-molecular-patterns (PAMPs), LPS is known for its ability to trigger the expression of inflammatory cytokines and the maturation of immune cells. The LPS-mediated signaling can be divided into MyD88-dependent and -independent (or TRIF-dependent) pathways(Lu *et al*, 2008) (Fig 2A). To identify the pathway that regulates *Slc29a3* expression upon LPS stimulation, we treated MyD88-deficient BMDMs with LPS or *E. coli* and monitored the mRNA level of *Slc29a3*. We found that both treatments effectively stimulated the upregulation of Slc29a3 in MyD88^-/-^ BMDMs, suggesting that MyD88 is dispensable for the LPS-induced Slc29a3 expression (Fig 2B). To examine if the MyD88-independent TRIF-mediated pathway regulates *Slc29a3* expression, we applied MRT 67307, a potent kinase inhibitor specifically blocking the function of TBK1 (TANK-binding kinase 1) and I-kappa-B kinase (IKK) epsilon (IKKε), together with LPS to WT or MyD88^-/-^ BMDMs. MRT67307 effectively blocked the LPS-induced *Slc29a3* expression and significantly inhibited the *E. coli* elicited responses in both WT (Fig 2C) and MyD88^-/-^ BMDMs (Fig 2D). These results suggest that the expression of *Slc29a3* is downstream of TRIF-mediated responses. TRIF-mediated signaling results in the activation of interferon regulatory factor 3 (IRF3), which enhances the production of interferon-β (IFN-β). The same dose and length of LPS treatment that upregulated *Slc29a3* expression effectively induced *IFNb1* production (Fig 2E) in BMDMs. To test if IFN-β modulates the transcription of *Slc29a3*, BMDMs were treated with IFN-β for different lengths of time. A clear time-dependent induction of *Slc29a3* by IFN-β stimulation was observed (Fig 2F), suggesting that LPS-induced *Slc29a3* expression is likely an IFN-β-mediated response.

**Figure 2.**
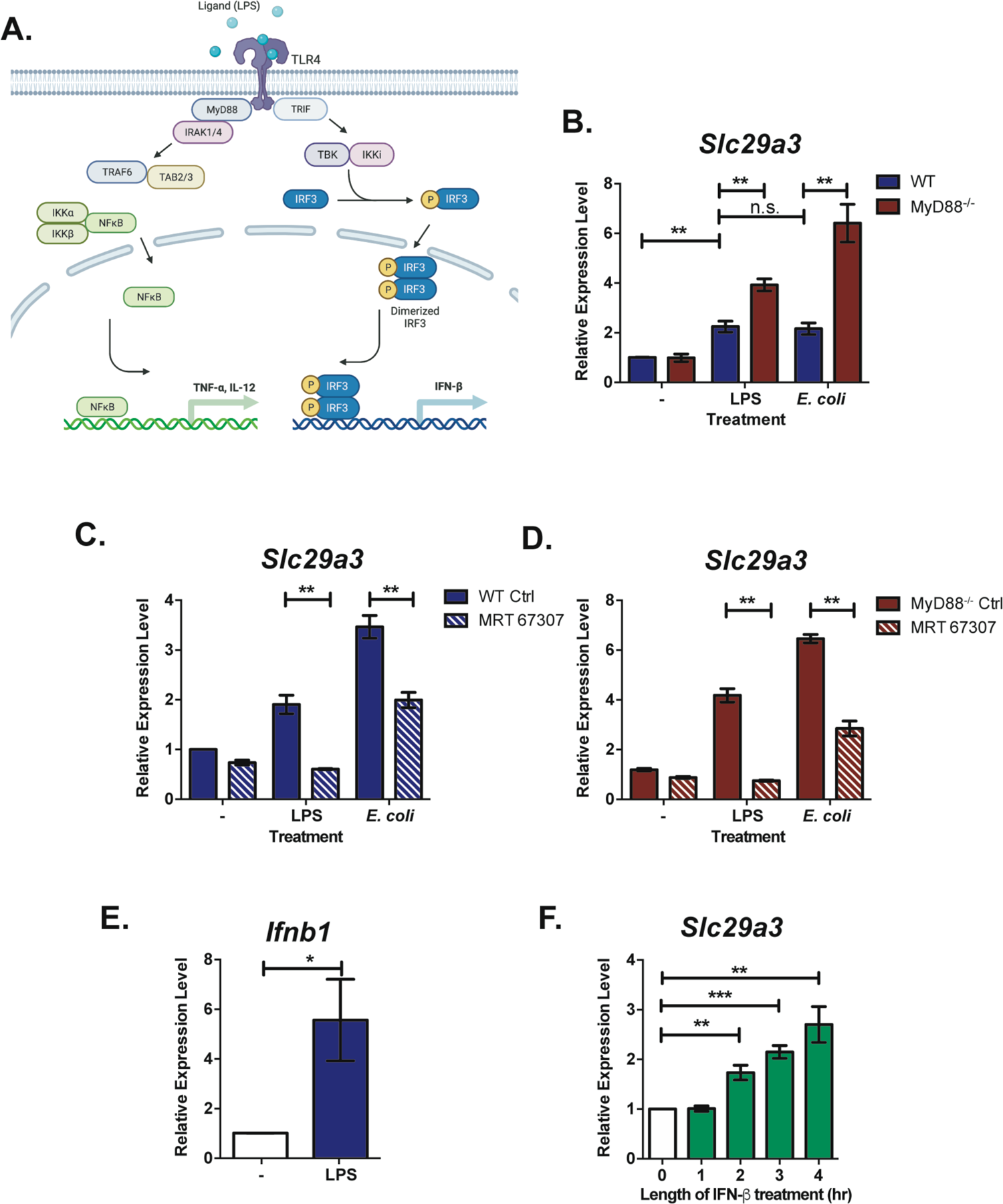
LPS induces ENT3 expression through the TLR4-TRIF axis. **(A)** The Schematic diagram of TLR4 signaling pathways. **(B)** BMDMs from WT or MyD88^-/-^ mice were treated with 10 ng/mL of LPS or live *E. coli* and subjected for *Slc29a3* expression analysis. **(C)** WT or **(D)** MyD88^-/-^ BMDMs were stimulated with LPS or *E. coli* in the presence of 10 μM TBK inhibitor, MRT 67307, and analyzed for *Slc29a3* expression. **(E)** The LPS-treated BMDMs were examined for *Ifnb1* expression. All BMDMs were treated with LPS or *E. coli* for 30 min, further incubated for a total of 6 hr and then harvested for RT-qPCR analyses. **(F)** BMDMs were treated with 100 U/mL of IFN-β for various length of time and subjected for *Slc29a3* expression. The results shown are combined from three independent experiments (N=3), unpaired two-tailed Student’s *t*-test was used for statistical analysis, \**p* < 0.05, \*\**p* < 0.01, \*\*\**p* < 0.001, n.s. no significance.

### IFN-β is the crucial mediator that activates the transcription of *Slc29a3*

IFN-β belongs to the type I IFN family, it mediates signaling events through the type I interferon receptor (IFNAR) complex to activate the transcription of ISGs (Platanias, 2005) (Fig 3A). To investigate the importance of the IFN-β-mediated signaling cascade in regulating *Slc29a3* transcription, we stimulated IFNAR^-/-^ BMDMs with LPS, live *E. coli*, or IFN-β. None of the stimuli could induce the upregulation of *Slc29a3* in IFNAR^-/-^ BMDMs, demonstrating that Slc29a3 expression is directly under the control of IFN-β (Fig 3B). Upon ligand binding, IFN-β initiates the activation of the JAK-STATs pathway and recruits IRF9 to form the IFN-stimulated gene factor 3 (ISGF3). The ISGF3 translocates to the nucleus leading to the transcription of ISGs(Wang *et al*, 2017). To further examine if Slc29a3 is an unidentified ISGs, we treated STAT1^-/-^ BMDMs with LPS, *E. coli*, or IFN-β but none of them could induce the expression of *Slc29a3* (Fig 3C). The IFN family is comprised of two major classes of related cytokines –type I and type II IFNs. Harboring antiviral activities, the only type II IFN, IFN-γ, does not have marked structural homology to type I IFNs and interacts with a different type II receptor (IFNGR)(Platanias, 2005). To test the effects of major IFNs on *Slc29a3* induction, we treated BMDMs with different doses of IFNs and profiled the expression of *Slc29a3*. We found that both type I IFNs, IFN-α (Fig 3C) and IFN-β (Fig 3D), are effective inducers for *Slc29a3* expression, and the induction peaked rapidly at 4 hours upon stimulation and remained above the resting level at least for 48 hours. On the other hand, IFN-γ elicited only moderate upregulation of *Slc29a3* at later time points (Fig 3E). These results suggest that the transcription of *Slc29a3* is likely a direct downstream target of type I IFN-IFNAR signaling event. Moreover, these findings strongly imply that the metabolite transporter, ENT3, might be an unrecognized ISG.

**Figure 3.**
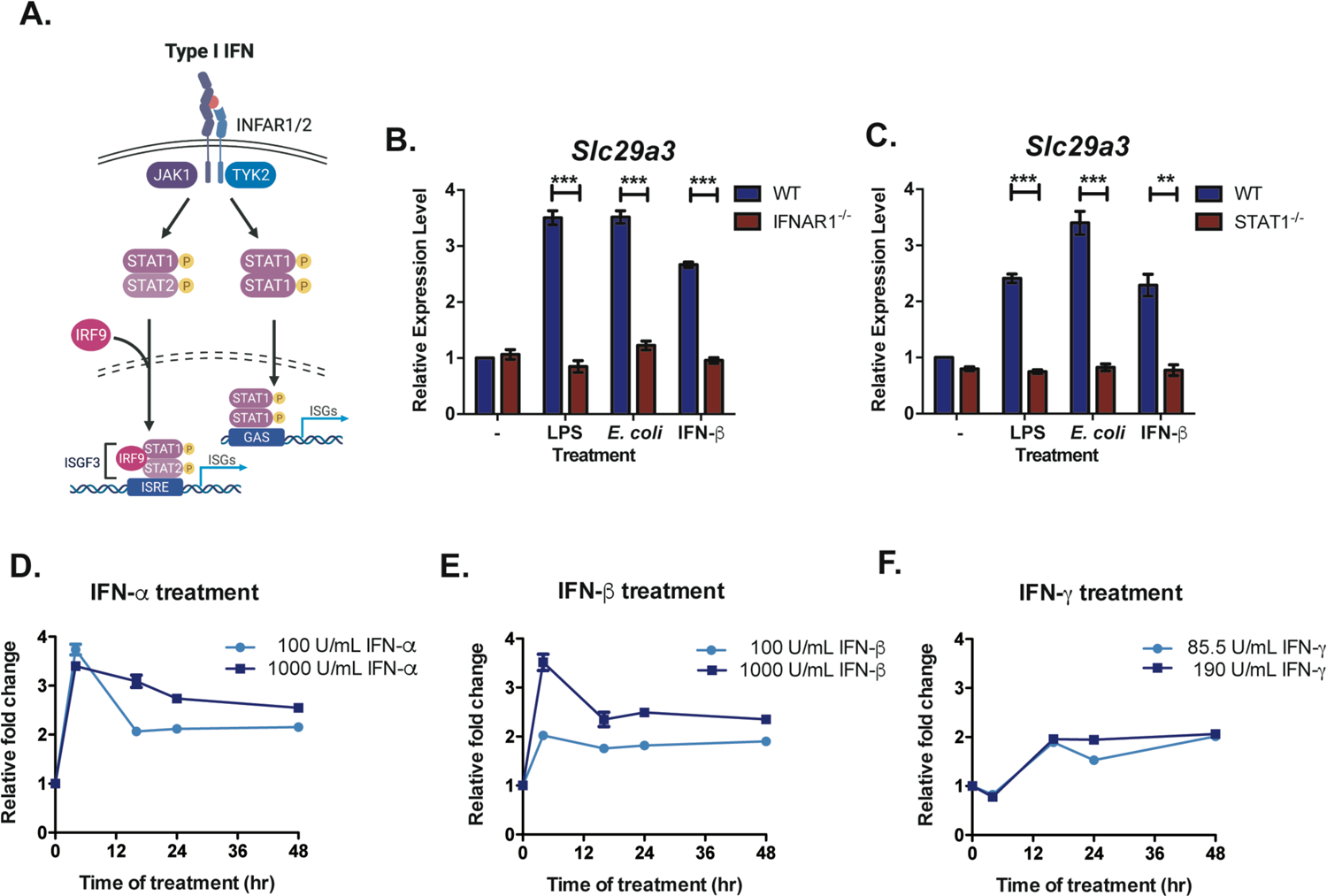
Type I IFN upregulates ENT3 transcripts through IFNAR1-STAT1 axis. **(A)** The brief schematic diagram of type I IFN downstream signaling cascades. **(B, C)** BMDMs from WT, IFNAR1^-/-^ **(B)** or STAT1^-/-^ **(C)** mice were treated with LPS and live *E. coli* or IFN-β, harvested and examined for *Slc29a3* induction. The expression dynamic of *Slc29a3* was profiled in BMDMs treated with two different doses of IFN-α **(D)**, IFN-β **(E)**, IFN-γ **(F)** for various length of time as indicated. **(B, C)** The results shown are combined from three independent experiments (N=3), unpaired two-tailed Student’s *t*-test was used for statistical analysis, *\*\*p* < 0.01, *\*\*\*p* < 0.001, n.s. no significance. **(D-F)** Shown is the representative data of two independent experiments. The expression level was calculated relative to *Rpl19* and normalized to untreated group at time 0 as 1.

### ENT3 is an IFN-stimulated gene (ISG)

Upon sensing the pathogen and the production of IFNs, hundreds of ISGs are transcribed to arm the infected and nearby cells to an antiviral state – including enhanced detection of pathogens and initiation of the pathogen removal machinery. Although a variety of genes comprises ISGs, they share the common goal of combating the pathogen and regaining homeostasis(Schneider *et al*, 2014). To explore the possibility that ENT3 is an ISG, we extracted the results from the microarray datasets curated in the INTERFEROME database (http://www.interferome.org/interferome/home.jspx). We found that Slc29a3 is upregulated to the same level as other validated positive regulators and antiviral effector ISGs in response to IFN-β treatment (Fig 4A), further supporting the notion that *Slc29a3* is one of the Interferon-regulated genes (IRGs). To investigate if a direct interaction between the IFNAR-STAT1 pathway and Slc29a3 expression at the transcriptional level exists, we searched the ChIP-Atlas database (http://chip-atlas.org/), which curated ChIP-seq data for potential STAT1 binding sites on Slc29a3 gene locus. We identified three potential STAT1 binding sites in the promoter and 3’ untranslated region (UTR) of Slc29a3 from the database search (Fig 4B). To confirm this result, we performed ChIP-qPCR on IFN-β treated BMDMs (Supplemental Fig 3), and clear STAT1 binding was found in the Slc29a3 promoter region (Fig 4E). In contrast, no statistically significant differences were found in the two additional potential STAT1 binding sites within the 3’ UTR compared with untreated control (Fig 4C, D). Together, these results support the idea that ENT3 is an ISG under the direct regulation of the IFN-β-STAT1 pathway.

**Figure 4.**
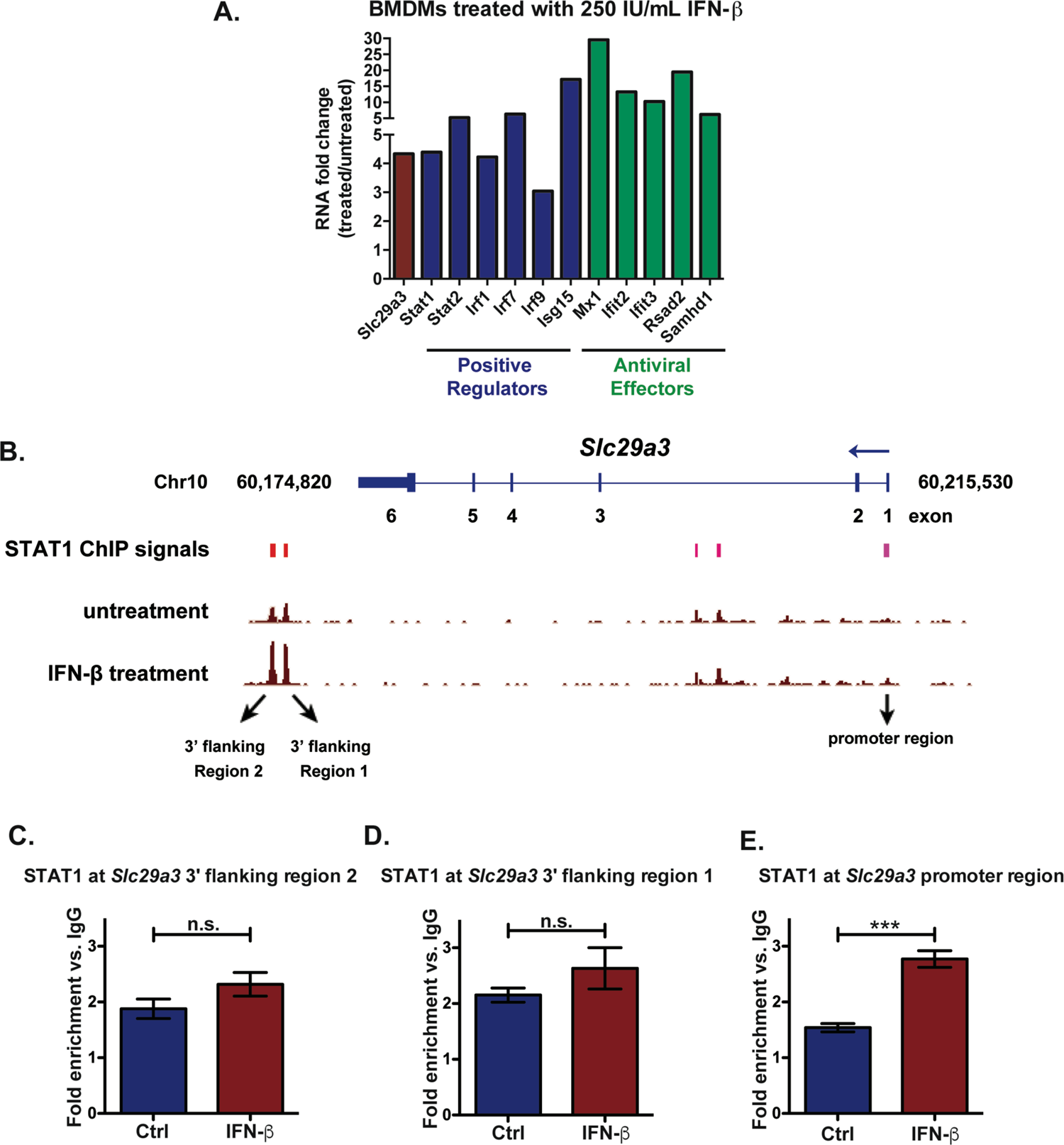
Slc29a3 has a STAT1 binding site on its promoter region. **(A)** The induction fold change (treated/untreated) of *Slc29a3* and selected ISG transcripts were curated from INTERFEROME database. Shown is data extracted from 250 IU/mL IFN-β treated BMDMs. **(B)** Potential STAT1 interacting sites on Slc29a3 were identified using the ChIP-Atlas database. The red bars marked the STAT1 binding sites while the height of the bars represented the frequency of STAT1 binding on the specified region. **(C-E)** Primers targeting 3’ flanking region 2 **(C)**, 3’ flanking region 1 **(D)**, and promoter region **(E)** were used to quantify the STAT1 binding in BMDMs upon 100 U/mL IFNβ 4 hr stimulation via qChIP assay. The results shown are combined from two independent experiments (N=2, n=4), unpaired two-tailed Student’s *t*-test was used for statistical analysis, *\*\*\*p* < 0.001, n.s. no significance.

### Macrophages upregulate ENT3 upon sensing viral infection, yet the deficiency of ENT3 ameliorates EMCV-induced pathology and lowers viral load *in vivo*

Since IFN-mediated innate immune response is a robust arsenal to fight off invading pathogens, especially viruses, we wondered if the viral infection could trigger the upregulation of ENT3. We chose two virus types to challenge the macrophages – encephalomyocarditis virus (EMCV) and enterovirus 71 (EV71). Both EMCV and EV71 are small non-enveloped RNA virus belonging to *Picornaviridae*. EMCV is known to cause myocarditis and encephalitis in mice, while EV71 infection is the common cause of human’s hand, foot, and mouth disease (HFMD). We observed EMCV infection increased the transcription of Slc29a3 in BMDMs (Fig 5A). THP-1 derived human macrophages also responded to the EV71 challenge by the upregulation of *Slc29a3* (Fig 5B). To confirm these observations, we challenged BMDMs with Poly I:C, the synthetic mimetic of a double-stranded RNA capable of stimulating the immune system through the TLR and RLR families that drives potent type I IFN responses. Poly I:C stimulation effectively induced the expression of Slc29a3 in BMDMs (Fig 5C). Together, these results demonstrate that the upregulation of ENT3 in macrophages is an early response to viral invasion. While it is well known that ISGs are synergistic in blocking virus entry and inhibiting viral translation and replication, the role of ENT3 as an ISG upon viral infection is unclear. To study the participation of ENT3 in antiviral response *in vivo*, Cx3cr1-cre ENT3^fl/fl^ mice were challenged with EMCV. To our surprise, the Cx3cr1-cre ENT3^fl/fl^ mice were more resistant to the EMCV-induced lethality and showed better survival (Fig 5D). In agreement with the survival curve, the Cx3cr1-cre ENT3^fl/fl^ mice exhibited clearly milder encephalitis phenotype (Fig 5E). We performed the plaque assay to measure the viral titer in the EMCV-infected mice and found significantly lower viral load in the Cx3cr1-cre ENT3^fl/fl^ mouse brains (Fig 5F and G). These results together demonstrated that although macrophages upregulate the expression of ENT3 upon viral challenge, the absence of ENT3 is unfavorable for viral replication.

**Figure 5.**
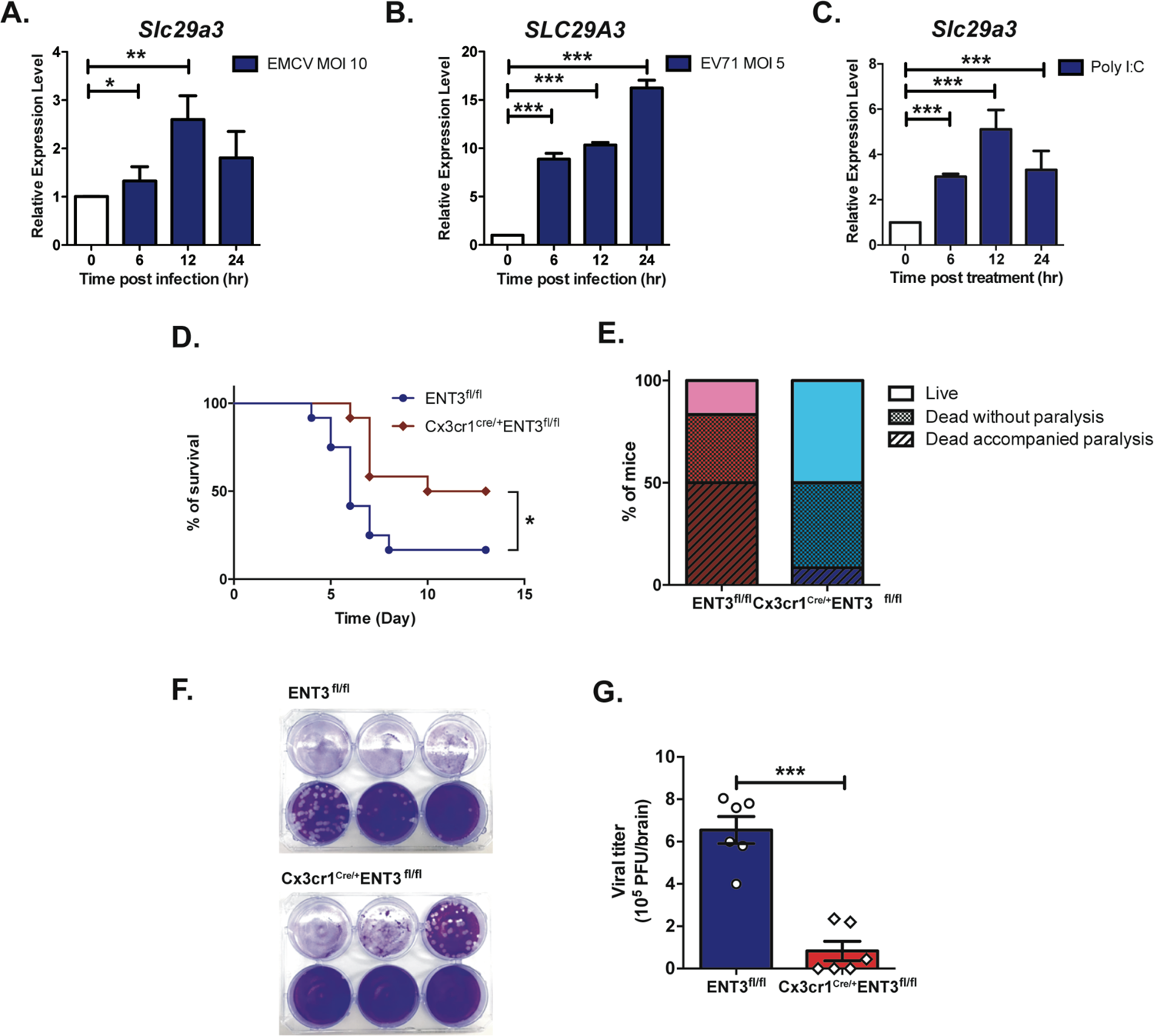
The expression of ENT3 is responsive to viral infection, yet Cx3cr1^cre/+^ENT3^fl/fl^ mice are resistant to EMCV infection. **(A)** BMDMs were infected with EMCV (MOI=10) and harvested at indicated time points. **(B)** THP-1 derived macrophages were challenged with EV71 (MOI=5) and collected at different time points. **(C)** BMDMs were co-cultured with 50 µg/mL poly (I:C) with various length of time and harvested. The mRNA expression of *Slc29a3* or *SLC29A3* was measured by RT-qPCR. The expression level was calculated relative to *Rpl19* or *RPLP0* and normalized to untreated group at time 0 as 1. Results are combined from three independent experiments (N=3), unpaired two-tailed Student’s *t*-test was used for statistical analysis, \**p* < 0.05, \*\**p* < 0.01, \*\*\**p* < 0.001. **(D, E)** ENT3^fl/fl^ (N=12) and Cx3cr1^cre/+^ENT3^fl/fl^ (N=9) mice were challenged with EMCV (10^4^ PFU, *i.p.*) and monitored for the survival **(D)** and disease severity **(E)** for two weeks. Survival results were analyzed by log-rank (Mantel-Cox) test, \**p* < 0.05. **(F)** ENT3^fl/fl^ and Cx3cr1^cre/+^ENT3^fl/fl^ mice (N=6) were challenged with 10^7^ PFU EMCV and measured the viral titer in the brain at post-infection day 3. Shown are representative plaque assay images. (G) EMCV viral titer in the infected mice. Unpaired two-tailed Student’s *t*-test was used for statistical analysis, \*\**p* < 0.01.

### The presence of ENT3 is vital for viral replication

The lower viral loads in Cx3cr1-cre ENT3^fl/fl^ mouse may have two possible explanations: the ENT3-deficient mouse had hyper-activated IFN responses that rapidly cleared the viruses, or the defect in ENT3 impacted viral replication. To probe these two possibilities, we infected BMDMs with EMCV *in vitro* and followed BMDMs’ IFN response. We found that upon EMCV infection, ENT3^-/-^ BMDMs had significantly lower IFN-β production than WT (Fig 6A), suggesting that ENT3^-/-^ BMDMs do not have a hyperactive antiviral response. To explore if this result was due to the altered cytosolic innate sensing in the absence of ENT3, we introduced poly I:C, the long synthetic analog of dsRNA, via liposome into BMDMs. We found a lower IFN-β expression upon liposome-mediated poly I:C challenge in ENT3^-/-^ BMDMs compared with WT, suggesting the cytosolic RNA helicases retinoic acid-inducible protein I (RIG-I) and melanoma differentiation-associate gene 5 (MDA-5) modules may be hypo-functional (Fig 6B). These results rule out the hyperactive IFN responses in ENT3^-/-^ macrophages instead argues for the role of ENT3 in EMCV replication. By measuring the transcription of EMCV viral protein 2A2B, we found the EMCV replication was almost completely abolished in ENT3^-/-^ BMDMs (Fig 6C). As metabolite transporter, we speculated that ENT3 might be an ISG involved with the regulation of macrophage cellular metabolism. Macrophages undergo metabolic reprogramming upon infection to support their effector functions; on the other hand, viruses also actively manipulate host cell metabolism to establish an optimal environment for their propagation. To evaluate if ENT3^-/-^ BMDMs are metabolically distinct from WT BMDMs, we applied the proton efflux assays to measure the glycolytic and mitochondrial activities. The results showed no differences in OCR (Fig 6D) yet significantly higher ECAR in ENT3^-/-^ BMDMs compared with WT (Fig 6E), suggesting that the abolished viral replication in ENT3^-/-^ BMDMs is unlikely due to defective glycolytic or mitochondrial activities of BMDMs. However, upon activation-induced metabolic reprogramming process, BMDMs defective in ENT3 function may have disturbances in metabolic pathways other than glycolysis. To explore this possibility, we treated WT and ENT3^-/-^ BMDMs with IFN-β and performed RNAseq analyses. Interestingly, but not unexpectedly, GSEA on KEGG pathways revealed the strong association of nucleotide-related activities, including DNA replication, nucleotide excision repair, and pyrimidine metabolism in ENT3^-/-^ BMDMs compared with the WT group (Fig 6F). The increased nucleotide excision repair and pyrimidine metabolism implied the disturbed nucleoside/nucleotide supply in ENT3^-/-^ BMDMs. These observations prompted us to hypothesize that ENT3 may have an essential role in providing increased pools of free nucleotides necessary for rapid viral genome replication. To test this possibility, we challenged ENT3^-/-^ BMDMs with EMCV and measured the viral replication in the absence or presence of ribonucleosides (rN) supplements. We noted that the addition of rN did salvage the viral replication in ENT3^-/-^ BMDMs in a dose-dependent fashion (Fig 6G); however, the rescue effect on viral replication is minute compared to WT BMDMs without rN addition. These results suggest that although ENT3 deficiency impacts the nucleotides pool to support viral replication, ENT3 likely has an additional crucial role in viral replication and survival.

**Figure 6.**
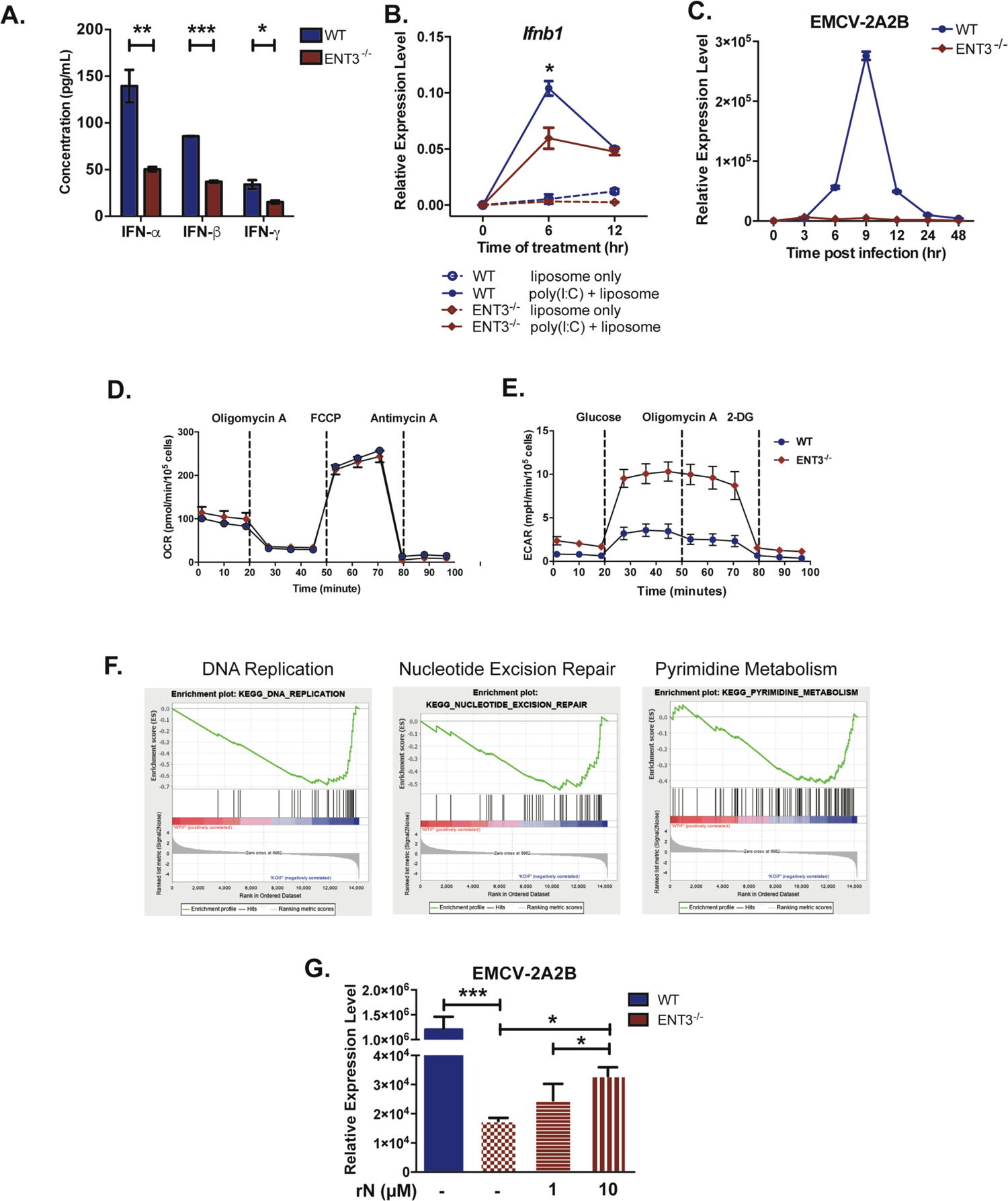
EMCV fails to replicate in ENT3^-/-^ BMDMs. **(A)** WT and ENT3^-/-^ BMDMs were infected with EMCV (MOI=10) and harvested at 24 hr post-infection to measure the production of IFN-α, IFN-β, and IFN-γ by ELISA (N=3). **(B)** WT and ENT3^-/-^ BMDMs were challenged with 1 µg/mL of liposome-packed poly (I:C) for the length of time indicated and harvested to evaluate the *Ifnb1* expression by RT-qPCR. Shown are combined results from three independent experiments (N=3). **(C)** WT and ENT3^-/-^ BMDMs were infected with EMCV (MOI=10) for 2 hr and allowed for viral replication for the various length of time as indicated. The expression of EMCV-2A2B was determined by RT-qPCR. Representative data from three independent experiments are shown. The relative gene expression was normalized to *Rpl19*, unpaired two-tailed Student’s *t*-test was used for statistical analysis, \*\*\**p* < 0.001. **(D, E)** The respiratory capacity **(D)** and glycolysis capacity **(E)** of WT and ENT3^-/-^ BMDMs were measured by extracellular flux analysis. Shown are the representative results of OCR and ECAR kinetics from two independent experiments. **(F)** GSEA of RNAseq results generated from WT and ENT3^-/-^ BMDMs treated with 1000 U/mL IFN-β for 4 hr. Shown are the significantly enriched pathways in ENT3^-/-^ BMDMs. **(G)** The replication of EMCV in ENT3^-/-^ BMDMs were evaluated in the presence of exogenous 1 or 10 µM of ribonucleosides (A, U, C, G) 9 hr post-infection. Shown are combined results from three independent experiments (N=3). unpaired two-tailed Student’s *t*-test was used for statistical analysis, \**p* < 0.05, \*\*\**p* < 0.001.

### ENT3 supports the release of viral RNA into the cytosol

Productive viral infection requires multiple steps: entry, genome replication, and exit (Ryu, 2017). On the other hand, the adequate clearance of the virus requires recognition, activation of IFN responses, and induction of adaptive immunity. To this point, we had tested the involvement of ENT3 in influencing the general metabolism of macrophage, intracellular nucleoside availability, or the IFN-mediated responses. However, none of them can explain the impact of ENT3 on viral replication. These results prompted us to speculate if the virus requires functional ENT3 for its entry into the cell. Viral entry can be divided into attachment, penetration, and uncoating. As a metabolite transporter, ENT3 translocates nucleosides and nucleotides through lysosomal membranes and contributes to maintaining lysosomal pH, implying its potential involvement in virus uncoating. To test this possibility, we labeled viral RNA with Syto82 dye (Hover *et al*, 2018) and allowed these viruses to infect target cells (Fig 7A). Comparing WT to ENT3^-/-^ BMDMs, we found that a substantial amount of EMCV viral RNA was trapped in the ENT3^-/-^ BMDMs (Fig 7B, yellow arrowheads) while most of the viral RNA had entered and dissipated into the cytoplasm in the WT BMDMs. These images were further quantified and revealed that a significantly higher percentage of ENT3^-/-^ BMDMs were Syto82 positive (Fig 7C) and were with stronger signals compared to the WT group (Fig 7D). These results support the notion that ENT3 has a vital role in the viral uncoating step. To further explore the importance of ENT3 in viral replication, we included an additional virus strain, SARS-CoV2, for testing. The ENT3 knockdown Calu-3 cells (human lung epithelial cells) were generated using the lentiviral-shRNA system (Fig 7E). To examine if the presence of ENT3 affects the replication of SARS-CoV2, the control, and ENT3 knockdown Calu-3 were infected with either SARS-CoV2 or delta variant (B.1.617.2) and evaluated. We found that even ∼ 50% of ENT3 knockdown significantly decreased the viral replication of both SARS-CoV2 (Fig 7F) and delta variant (Fig 7G) judging by the viral RNA amount and plaque assay (Supplemental Fig 4). These results suggest ENT3 is a key component in viral propagation and imply that modulating ENT3 expression could have therapeutic applications.

**Figure 7.**
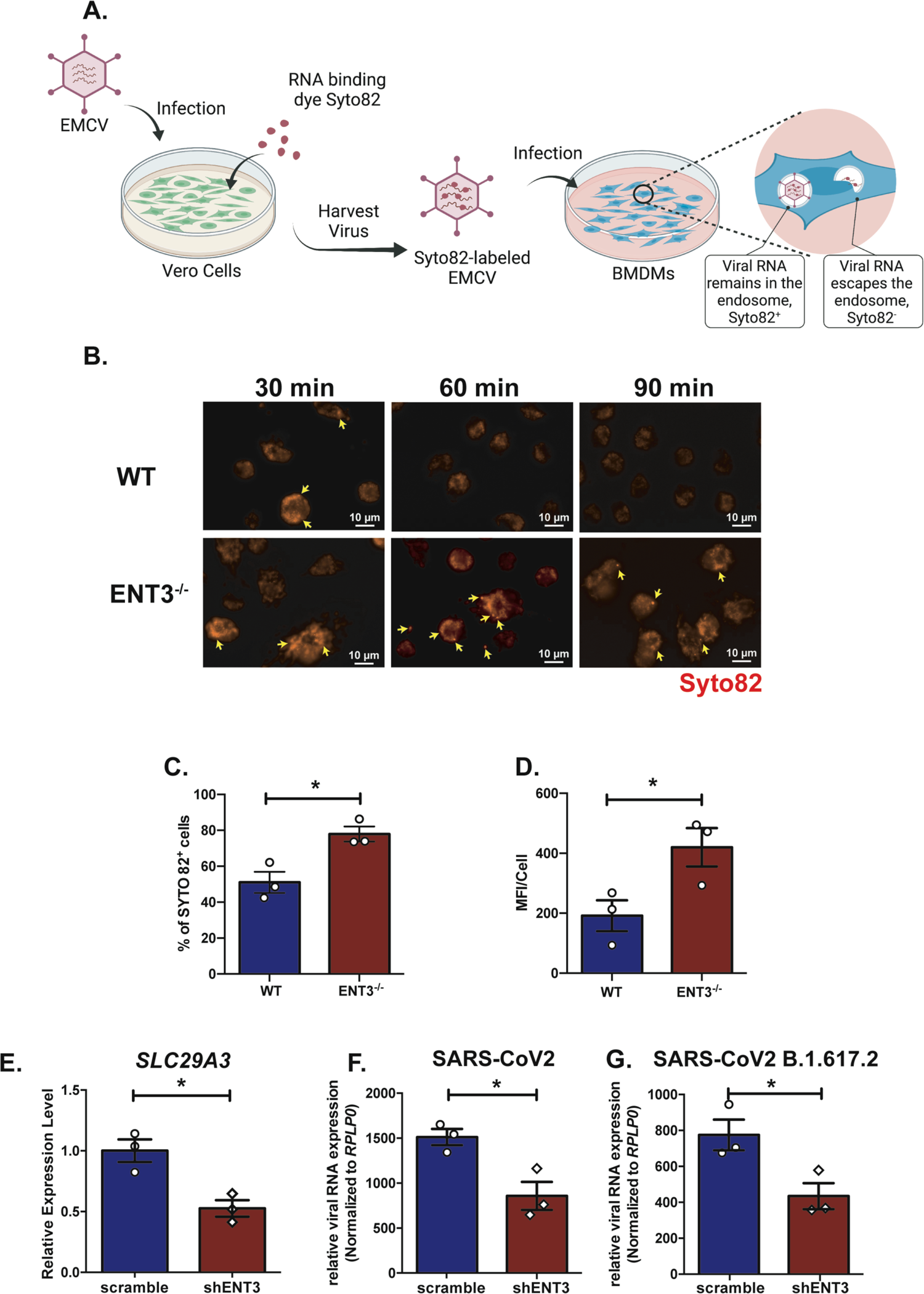
ENT3 supports the uncoating of virus, and the replication of SARS-CoV-2 requires functional ENT3. **(A)** Illustration of strategy to track the viral uncoating upon infection. **(B)** WT and ENT3^-/-^ BMDMs were infected with Syto82-labeled EMCV (MOI=1) and subjected to live imagining at 30-, 60- and 90-min post-infection. The arrowheads indicated the Syto82^+^ virus puncta in the cell. **(C, D)** Quantification of Syto82 signal in BMDMs at 60 min time point was performed by enumerating Syto82^+^ cells (%) **(C)** and the mean fluorescent intensity (MFI) per Syto82^+^ cell **(D)**. Results were generated images from 3 independent experiments, each with 3 to 4 fields analyzed. Unpaired two-tailed Student’s *t*-test was used for statistical analysis, \**p* < 0.05. **(E)** ENT3 knockdown in Calu-3 cells was performed by lentiviral shRNA system. The knockdown efficiency was confirmed by RT-qPCR. The *SLC29A3* expression level was calculated relative to *RPLP0* and normalized to scramble control as 1. **(F, G)** ENT3 knockdown Calu-3 cells were infected with SARS-CoV-2 original strain **(F)** or delta variant **(G)** at MOI=0.1 for 1 hr, and subjected for the viral replication analysis at 24 hr post-infection by RT-qPCR. The SARS-CoV-2 E gene expression level was calculated relative to *RPLP0* to normalize the cell number. Shown are combined results from three independent experiments (N=3). Unpaired two-tailed Student’s *t*-test was used for statistical analysis, \**p* < 0.05.

## Discussion

This study unveils ENT3 as an early metabolic responder upon bacterial and viral challenge. Through dissecting the signaling events regulating the expression of ENT3, we also reveal that ENT3 is an ISG. Intriguingly, it is an ISG being taken advantage of by a virus to facilitate its replication. These results demonstrate that metabolic preparedness is a vital part of innate immunity and how pathogens have co-evolved to exploit this machinery to their benefits. These observations may provide a novel path to develop therapeutic interventions against viral infection.

Our understanding of the intimate connection between cell metabolism and function has recently progressed rapidly. We now know that immune effector functions, such as cytokine production or differentiation, are directly coupled to specific metabolic changes in the cell and can be affected by multiple metabolic pathways. For example, metabolic remodeling is tightly linked to macrophages’ development, activation, function, and differentiation (O’Neill & Pearce, 2015). We observed a significant increase of glycolysis in ENT3^-/-^ BMDMs. It is possible that ENT3 participates in the energy metabolism or the metabolic alteration compensates for ENT3 deficiency. It is known that the increase of glycolysis leads to the accumulation of acetyl-CoA, which is associated with histone acetylation and the relaxed chromatin state, and positive transcription regulation in macrophages(Kelly & O’Neill, 2015). It has been shown that the glycolytic enzyme, hexokinase, not only can act as a pattern recognition receptor (PRR) but also, together with mTORC1, plays a critical role for the NOD-like receptor family pyrin domain containing 3 (NLRP3) inflammasome assembly in macrophages(Moon *et al*, 2015). The conserved RNA-enzyme-metabolite (REM) networks, in which the metabolic enzymes bind mRNA and regulate proteins’ expression, also serve as additional measures translating metabolic status to immune cell function(Chang & Pearce, 2016). No noticeable inflammatory signs are reported on ENT3^-/-^ macrophages or mice(Hsu *et al*, 2012), implying that the disturbed balance between glycolysis and oxidative phosphorylation does not directly impact the activation status of ENT3^-/-^ cells but instead is likely to compensate the metabolic needs. Following the glycolytic flux in ENT3^-/-^ cells may gain further insights into how ENT3 supports cellular metabolism.

While the growing literature underscores that the metabolic program of immune cells can lead to changes in their effector functions, the signaling events leading to the expression of metabolite transporters support the metabolic preparedness or the types of metabolite transporters responding to different pathogen challenges are still underexplored. In this study, we found that type I IFN-mediated signaling directly regulates the expression of ENT3. It will be of great interest to identify the metabolite transporters responding to IFN stimulation and gain an overview of the metabolic module responding to viral infection. Moreover, it is plausible that different types of stress, such as damage-associated molecular patterns (DAMP), intracellular or extracellular microbes, can induce the expression of distinct metabolite transporter clusters.

Although the observation that ENT3 supports viral replication is exciting, a clear limitation of this study is that the cellular function of ENT3 upon infection is yet to be clarified. Since the expression of ENT3 is under the direct regulation of IFN-β, it is likely a part of the antiviral program. The suppressed viral replication in ENT3 knockdown or knockout cells may have two possible explanations. One is that the virus fails to enter the cytoplasm; the other is that the ENT3 defective cells cannot sustain viral replication. As the supplement of the nucleosides *in vitro* cannot overcome the defective viral replication in ENT3^-/-^ cells, the result suggests the defect lies in the viral genome release step. Sequential and proper acidification is crucial for viral entry into the cytoplasm(Mudhakir & Harashima, 2009), which makes the known phenotype of lysosomal pH elevation in ENT3^-/-^ cells(Hsu *et al*, 2012) a reasonable explanation for the inefficient or defective viral uncoating in the absence of ENT3. Experimentally, the lack of ENT3-specific functional inhibitors makes it unachievable to dissect the roles of ENT3 on lysosomal nucleoside transportation and pH regulation – because ENT3 is a pH-dependent transporter and the alteration of pH value directly affects ENT3 function. Moreover, the disturbed lysosomal function can have various impacts on cellular metabolism in addition to nucleoside availability, and we do not exclude the involvement of effects secondary to ENT3 deficiency.

As nucleoside transporter has a vital part in salvage pathway, we consider ENT3 may also take part in fulfilling the nucleotide metabolic demand of immune responses. Nucleosides are the essential components of DNA, RNA, and cofactor of the metabolic enzyme such as nicotinamide adenine dinucleotide (NAD)(Suganuma & Workman, 2021). It is plausible that ENT3 contributes to the transcription or even epigenetic regulation upon activation. We did observe lower transcription of *Ifnb1* upon poly I:C transfection in ENT3^-/-^ BMDMs compared to WT, which argues the possibility that ENT3 participates in RNA transcription. Whether ENT3 contributes to global RNA transcription or supplements the increased nucleoside demands upon stimulation still need further investigation.

As an ISG, it is almost anti-intuitive that ENT3 supports viral replication. However, microbes have evolved mechanisms altering immune cell metabolism to their survival benefits. For example, *Staphylococcus aureus* targets the ENT1-mediated purine salvage pathway to induce the intoxication of phagocytes(Winstel *et al*, 2018). *Trypanosome cruzi* preferentially targets pancreatic beta cells and results in the insulinemia accompanied with hyperglycemia of the host(Dufurrena *et al*, 2017) to dampen the immune responses. At a cellular or systemic level, the pathogens modulate host immune response via metabolic perturbance to ultimately prolong the infection and enhance the transmissibility. On the other hand, tumor cells compete for glucose consumption and metabolically restrict T cell activity(Chang *et al*, 2015). The lactate released from these highly glycolytic tumor cells further polarizes tumor-associated macrophages into immunosuppressive phenotype(Vitale *et al*, 2019). Together, the tumor cells create a favorable microenvironment subverting immune responses via manipulating the metabolism. While many aspects of immune cell metabolic remodeling in different physiological settings are still not fully understood, there is a growing realization that inappropriate metabolic remodeling underlies many aberrant immune responses, and cellular metabolism manipulation can beneficially enhance or temper immunity. Here we demonstrate that the genetic knockout of ENT3 in macrophage confers the survival advantage upon lethal dose EMCV challenge in a murine model, and the downregulation of ENT3 is sufficient to decrease the replication SARS-CoV2 *in vitro*. Together, the results from this study identify that the expression of metabolite transporter can shape the outcome of infection and imply the potential of modulating ENT3 or other metabolite transporters’ expression as antiviral therapeutic means.

## Materials and Methods

### Mice

Wild type C57BL/6J mice were obtained from National Laboratory Animal Center, Taiwan. ENT3^-/-^ mice (Tang *et al*, 2010) were obtained from the Mutant Mouse Regional Resource Center. ENT3^fl/fl^ mice were generated from the National Research Program for Biopharmaceuticals, Taiwan. Cx3cr1^cre/+^ (B6J.B6N(Cg)-Cx3cr1^tm1.1(cre)Jung^/J) mice were acquired from Jackson Laboratory (Strain #:025524). Mice were housed and bred in a pathogen-free facility at the Laboratory Animal Center of National Yang Ming Chiao Tung University. MyD88^-/-^ mice (Hou *et al*, 2008) were provided by Dr. Guann-Yi Yu (NHRI). IFNAR1^-/-^ (Chen *et al*, 2013) and STAT1^-/-^ (Durbin *et al*, 1996) mice were provided by Dr. Chien-Kuo Lee (NTU). All experimental procedures were approved by the NYCU Institutional Animal Care and Use Committee.

### Cell lines

All cells were cultured at 37°C with 5% CO_2_ in a humidified incubator. Calu-3 cells, 293T cells, RD cells, and VeroE6 cells were cultured in complete DMEM comprising DMEM (Gibco), 2 mM L-glutamine (Gibco), 1 mM sodium pyruvate (Gibco), 1% Penicillin/Streptomycin (Gibco), 1% non-essential amino acids (Gibco), and 10% heat-inactivated fetal bovine serum (Gibco). THP-1 cells were cultured in RPMI (Gibco) supplemented with 2 mM L-glutamine, 1 mM sodium pyruvate, 1% Penicillin/Streptomycin, 1% non-essential amino acids, and 10% heat-inactivated fetal bovine serum. THP-1 cells were stimulated with 50 nM PMA (Sigma-Aldrich) for 2 hr and then washed with fresh media to induce the differentiation into macrophages. The activated THP-1 cells were further incubated for 24 hr, and the THP-1 derived macrophages were subjected for experiments.

### Bone marrow-derived macrophages

Total bone marrow cells were flushed out from the femur and tibia of 5- to 8-week-old mice and treated with ACK buffer to remove erythrocytes. Cells were cultured in complete DMEM at 37°C, 5% CO_2_ for 1 hr. Non-adherent cells were collected and cultured in complete DMEM supplemented with 10% L929 conditional medium at 37°C, 5% CO_2_ for 6 days. The media were replenished every 3 days. Mature BMDMs were harvested by 5 mM EDTA and rested at 37°C, 5% CO_2_ overnight for before experiments.

### Bacteria

*Escherichia coli* DH5α (Protech) were transfected with pUC19 plasmid which has ampicillin resistance gene as a selection marker. The titer of *E. coli* was determined by overnight culturing on 50 µg/mL ampicillin-contained LB-agar plates. *Escherichia coli* BL21-GFP provided by Dr. Jie-Rong Huang (NYCU) were recovered in 50 µg/mL ampicillin-contained LB-broth overnight and then induced GFP expression with 1 mM IPTG for 3 hr. The titer of *E. coli* BL21 was determined by OD_600_ (OD_600_ = 1 = 1 × 10^8^ CFU/mL).

### Viruses

EMCV (ATCC VR-1479) was amplified in Vero cells; EV-A71 genotype C strain (4643/Taiwan/1998) was amplified in RD cells. The viral titer was determined by plaque assay. Two clinical SARS-CoV-2 strains: hCoV-19/Taiwan/NTU13/2020 (SARS-CoV-2 original strain) and hCoV-19/Taiwan/NTU92/2021 (SARS-CoV-2 delta variant) isolated from sputum specimens of SARS-CoV-2-infected patients were propagated in VeroE6 cells, and the viral titer was determined by plaque assay.

## METHOD DETAILS

### *In vitro* bacterial challenge assays

For live *E. coli* preparation, *E. coli* were re-suspended in antibiotic-free complete DMEM; For heat-killed *E. coli* preparation, *E. coli* were incubated at 60°C for 60 min and then centrifuged, re-suspended in antibiotic-free complete DMEM; For gentamicin-killed *E. coli* preparation, *E. coli* were re-suspended in LB-broth containing 50 µg/mL gentamicin (Bio Basic) and shaken overnight at 37°C, re-suspended in antibiotic-free complete DMEM; For *E. coli* metabolites preparation, *E. coli* were re-suspended in antibiotic-free complete DMEM at 37°C for 30 min, and then the supernatants were harvested after filtration through 0.22 µm filter. The killing efficiency by heat and gentamicin was confirmed by overnight plating on ampicillin-free LB-agar plates. BMDMs were infected with MOI 10 or MOI 100 *E. coli* in antibiotic-free complete DMEM and incubated at 37°C, 5% CO_2_ for 30 min. Subsequently, BMDMs were washed with warm PBS to remove non-internalized *E. coli* and incubated in penicillin/streptomycin-contained complete DMEM at 37°C, 5% CO_2_ to the indicated time point. For phagocytosis inhibition or TRIF-pathway inhibition assay, BMDMs were pre-treated with 10 µM cytochalasin D (Invitrogen) or 10 µM MRT67307 (Invivogen) for 1 hr, and allowed to co-incubate with the treatments. For Cytochalasin D inhibition efficiency test, GFP-expressing *E. coli* BL21 were pre-stained with pHrodo Red (Invitrogen) in PBS (10^8^ CFU/2 mL) at room temperature for 30 min and then infected BMDMs at MOI 100 in the presence or absence of Cytochalasin D at 37°C, 5% CO_2_ for 30 min. Images were captured by Zeiss Axio Observer 7 40X lens after washing out the extracellular *E. coli* with warm PBS.

### *In vitro* viral infection assays

BMDMs were infected with MOI 10 EMCV in 2% FBS complete DMEM at 37°C, 5% CO_2_ for 2 hr. Subsequently, BMDMs were washed with warm PBS to remove non-attached EMCV and incubated in 10% FBS complete DMEM at 37°C, 5% CO_2_ for various length as indicated in the figures. For rN supplementation test, different doses of exogenous ribonucleosides (A, U, C, G) were added into culture medium after washing out the non-attached EMCV. THP-1 derived macrophages were infected with MOI 5 EV71 in 2% FBS complete RPMI at 37°C, 5% CO_2_ for 1 hr. After viral adsorption, THP-1 derived macrophages were washed with warm PBS to remove non-attached EV71 and incubated in 10% FBS complete RPMI at 37°C, 5% CO_2_ to the indicated time point. Calu-3 cells were infected with MOI 0.1 SARS-CoV-2 in DMEM supplemented with 2% TPCK-trypsin for 1 hr at 37°C, 5% CO_2_. The virus-containing supernatants were then removed and the infected cells were further cultured in 2% FBS complete DMEM for 24 hr before analysis.

### Plaque assay

For EMCV infection, viral titer quantification was performed on Vero cells. Culture supernatants or brain tissue samples were added to the monolayer Vero cells at 37°C, 5% CO_2_ for 2 hr. The virus-containing supernatants were removed, and cells were maintained in 1% agarose (Lonza) overlay medium. After 30 hr incubation, cells were fixed with 4% paraformaldehyde for 30 min and then stained with 0.5% crystal violet for 30 min to visualize the plaque. For SARS-CoV-2 experiments, viral titer determination was performed as previously described (Cheng *et al*, 2021). Briefly, culture supernatants were treated to the monolayer VeroE6 cells for 1 hr at 37°C, 5% CO_2_. The virus-containing supernatants were removed, then cells were maintained in complete DMEM supplemented with 1% methylcellulose (Sigma-Aldrich) for 5 – 7 days. Finally, cells were fixed with 10% formaldehyde overnight and stained with 0.5% crystal violet for plaque visualization.

### Lentivirus-mediated ENT3 knockdown in Calu-3 cells

The shRNA lentiviral vectors targeting human ENT3 and a non-targeting control, scrambled shRNA, were purchased from National RNAi Core Facility Plateform at Institute of Molecular Biology, Academia Sinica, Taiwan. Each target lentiviral vector was co-transfected with psPAX2 packaging (Addgene) and pMD2.G plasmids (Addgene) into 293T cells to generate lentivirus. The lentiviral particles-containing supernatant for transduction were harvested and filtered through 0.45 µm filters, stored at −80°C. To generate ENT3 knockdown or control cells, Calu-3 cells were transduced with lentiviral particles-containing supernatant via spin-infection at 1,200 x g, 37°C for 2 hr in the presence of 8 µg/mL polybrene (Sigma-Aldrich). Subsequently, cells were recovered in antibiotic-free complete DMEM for 24 hr without puromycin selection and were ready for further experiments.

### *In vitro* TLR agonists and IFN stimulation

For PRR stimulation, BMDMs were stimulated with different doses of LPS (Invivogen) for 30 min, washed, and further incubated for 5.5 hr before harvest. BMDMs were transfected with 1 µg/mL poly(I:C) (Invivogen) packaged by Lipofectamine 2000 (Invitrogen) to activate intracellular PRRs via different entry routes. For cytokine stimulation, BMDMs were treated with different doses of IFN-α (BioLegend), IFN-β (R&D systems), and IFN-γ (BioLegend) for various length of time as indicated in the figures.

### RNA extraction and RT-qPCR

Total RNA was extracted by using TRI reagent (Invitrogen) following manufacturer’s protocol. Reverse transcription was performed by using SuperScript III Reverse Transcriptase (Invitrogen) with oligo (dT)_20_ (Invitrogen), dNTP mix (Invitrogen), and RNaseOUT (Invitrogen). qPCR was performed on StepOnePlus™ Real-Time PCR System (Applied Biosystems) by using Taqman universal master mix (Applied Biosystems) with Taqman Gene Expression Assay probe and primers (Thermo Fisher Scientific) or using iQ SYBR Green Supermix (Bio-Rad) with the primers listed in Table S1. The relative gene expression was normalized to *Rpl19* or *RPLP0* by using the 2-ΔCt method. The fold change of gene expression was normalized to unstimulated BMDMs (defined as 1) by using the 2-ΔΔCt method.

### Quantitative chromatin immunoprecipitation (qChIP)

ChIP assay was performed with SimpleChIP^®^ Enzymatic Chromatin IP Kit (Cell Signaling Technology). In brief, BMDMs treated with 100 U/mL IFN-β for 4 hr were fixed in 1 % formaldehyde for 10 min. Chromatin was fragmented by using micrococcal nuclease (Cell Signaling Technology), and the cell membrane was broken down by Bioruptor (Diagenode). 2% of samples were aliquoted for input controls, and the remaining samples were incubated with anti-STAT1 antibody (Cell Signaling Technology) and control antibodies (Cell signaling Technology) overnight at 4°C with rotation. The precipitated DNA was purified and conducted qPCR assay by using iQ SYBR Green Supermix with the primers listed in Table S1. The data were analyzed by the following formula: % of input = input dilution factor/bound dilution factor) × 2(input CT−bound CT) × 100 and then normalized to IgG control as 1.

### Measurement of IFNs by ELISA

Supernatants from cultured BMDMs were collected at 24 hr post-EMCV infection. Culture supernatants or serum samples were centrifuged at 10,000 x rpm, 4°C for 5 min before assay. IFN-α (Abcam), IFN-β (R&D systems), and IFN-γ (BioLegend) were assessed by ELISA kits according to the protocol of the manufacturer.

### Preparation of SYTO-82 labeled-EMCV

EMCV was labeled with SYTO-82 as described previously (Brandenburg *et al*, 2007). Briefly, Vero cells were infected with MOI 1 EMCV for 2 hr and then incubated with 25 µM SYTO-82 (Invitrogen) at 37°C, 5% CO_2_ for 4 hr. At end of the reaction, the cells were rinsed, harvested and freeze-thawed twice. Cell debris was removed by filtration through 0.45 µm filter. The viral titer was determined by plaque assay. BMDMs were treated with MOI 1 SYTO-82 labeled-EMCV for 10 min, rinsed and cultured at 37°C, 5% CO_2_ for 30, 60, or 90 min before subjected for observation. SYTO-82 signals were acquired on Zeiss Axio Observer 7. The images were processed by using the Zeiss Zen 2012 software and analyzed by using ImageJ.

### *In vivo* EMCV infection studies

9- to 13-week-old male Cx3cr1^cre/+^ENT3^fl/fl^ and ENT3^fl/fl^ mice were infected with 10^4^ PFU EMCV via *i.p.* for survival studies. The lethality and the disease phenotypes were closely monitored for two weeks. To determine the viral replication and acute cytokine responses *in vivo*, 10- to 11-week-old male mice were *i.p.* challenged with 10^7^ PFU EMCV and sacrificed 3 days post-infection. For the viral load examination in brains, brain tissues were homogenized in 1 mL PBS with plastic micro pestles, the supernatants were clarified by centrifugation twice at 3,000 rpm, 4°C for 10 min, and the viral titer quantified by plaque assay on Vero cells.

### Extracellular flux metabolic analysis

Oxygen Consumption Rate (OCR) and Extracellular Acidification Rate (ECAR) were recorded using the Seahorse XFe24 system (Agilent Technologies), and all procedures were performed following the manufacturer’s protocol. Briefly, BMDMs from WT or ENT3^-/-^ mice were seeded into XF24 culture microplate (Agilent Technologies) and incubated at 37°C, 5% CO_2_ overnight. Culture media were then replaced with complete DMEM without sodium pyruvate to evaluate the respiratory capacity of cells; and the complete DMEM without sodium pyruvate and glucose to measure the glycolysis capacity. After basal OCR or ECAR measurements, sequential injection of 1 µM oligomycin A (Sigma-Aldrich), 2 µM FCCP (Sigma-Aldrich), and 0.5 µM Antimycin A (Sigma-Aldrich) were applied for respiratory capacity measurements; 10 mM glucose, 1 µM oligomycin A, and 50 mM 2-DG (Sigma-Aldrich) were used for glycolysis capacity measurements. The parameters were recorded 3 times after each drug projection.

### RNA Sequencing and Gene Set Enrichment Analysis

BMDMs from WT or ENT3^-/-^ mice were treated with 1000 U/mL IFN-β for 4 hr, and the total RNA was extracted as described above. The RNA concentration was measured by Qubit fluorometer with Quant-iT RNA BR assay kit. The RNA integrity was checked by Agilent 2100 Bioanalyzer with RNA nanochip, and the RIN score of input samples had a minimum RIN score of 9. The RNA-sequencing libraries were prepared with Illumina TruSeq Stranded mRNA Sample Prep Kit and then sequenced on the NextSeq 550 platform with Illumina NextSeq High Output Kit v2.5 (75 cycles). The raw 75-bp reads were aligned to the *Mus musculus* genome assembly GRCm38 (mm10) and analyzed following the CLC Genomics Workbench 20.0.4 pipeline. Datasets were analyzed by using GSEA software (4.0.3) under KEGG pathways after the differential gene expression analysis which was calculated by using R package DESeq2 (1.30.0).

## QUANTIFICATION AND STATISTICAL ANALYSIS

Data analysis was performed by using Prism 6 (GraphPad) software. Results were analyzed by unpaired two-tailed Student’s *t*-test. Survival studies were analyzed by log-rank (Mantel-Cox) test. Data are presented as the mean ± SEM. Illustrations were made using BioRender.

## Acknowledgements

We would like to thank Dr. Lih-Hwa Hwang (NYCU) and Dr. Li-Chung Hsu (NTU) for sharing EV71 and EMCV, and You-Shen Lin (NTU) who generously provided technical supports for the EMCV-related experiments. Calu-3 cells were kindly provided by Dr. Szu-Ting Chen (NYCU). *E. coli* BL21-GFP were generously shared by Dr. Jie-Rong Huang (NYCU). We appreciate the insightful discussion with Dr. Chi-Ju Chen (NYCU). The RNAseq experiment was supported and performed at Genomics Center for Clinical and Biotechnological Applications of NCFB. We would like to acknowledge the services provided by the Biosafety Level-3 Laboratory of the First Core Laboratory from NTU College of Medicine and the Biosafety Level-3 Laboratory from NTUH, supported by grants from the Ministry of Science and Technology, Taiwan MOST-110-2740-B-002-006, MOST109-2327-B-002-009. This work is funded by MOST 108-2628-B-010-005, 109-2628-B-010-016, 110-2628-B-A49A-510 and Cancer Progression Research Center NYCU, from the Higher Education Sprout Project by MOE to C.-L. Hsu.

## Author Contributions

Y.-T.H., T.-L.T, C.-C.T., S.-Y.H., J.-W.H., and P.-Y.T. performed the experiments. T.-L.C., Y.-M.T., and S.-Y.C. executed and supported the SARS-CoV2 infection studies; P.-C.C. supported the ChIP-qPCR assay; C.-K.L. and G.-Y.Y. supported and provided multiple knockout mouse strains in the study. I.L.D., and C.-L.H. conceptualized the study, designed and supervised the experiments. Y.-T.H., and C.-L.H. analyzed and interpreted the data, and wrote the manuscript.

## Expanded View Figure legends

**Figure S1.**
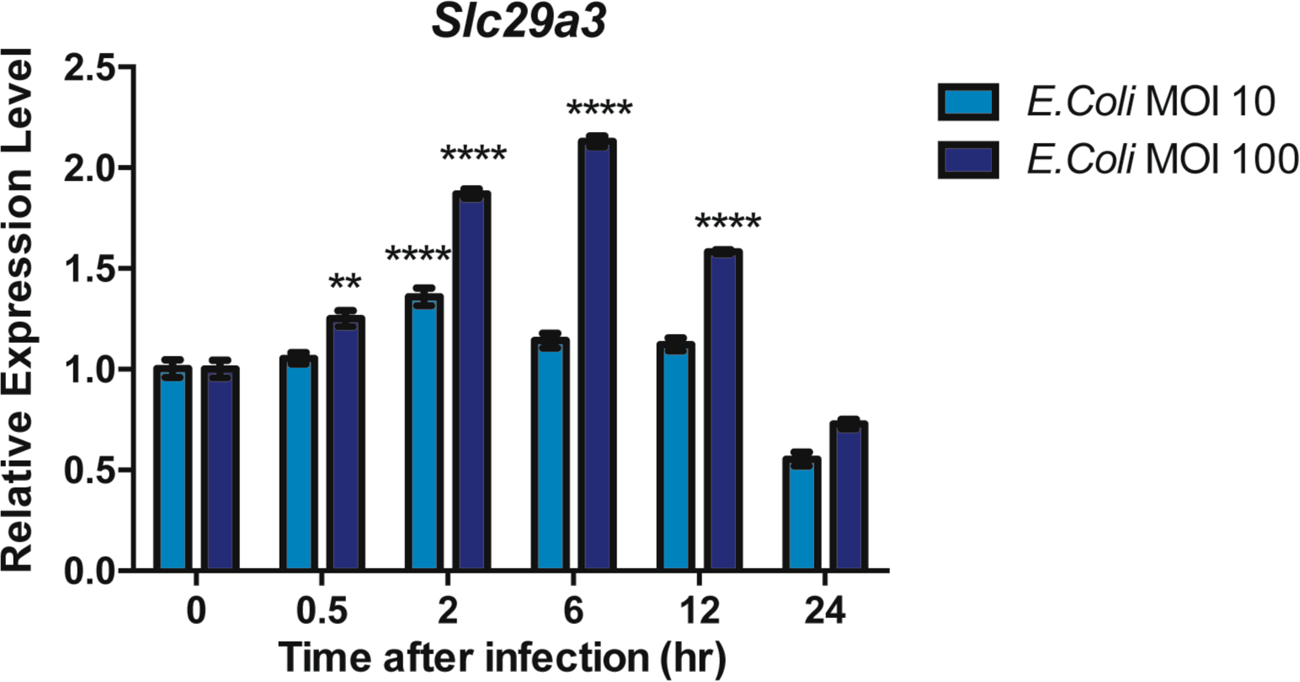
The transcription of *Slc29a3* in macrophages was upregulated during high-dose *E. coli* infection. BMDMs were treated with different MOIs of *E. coli* DH5α for 30 min and further cultured for various length of time as indicated. The dynamic of ENT3 expression was determined by measuring *Slc29a3* transcripts via RT-qPCR. Representative data from three independent experiments are shown.

**Figure S2.**
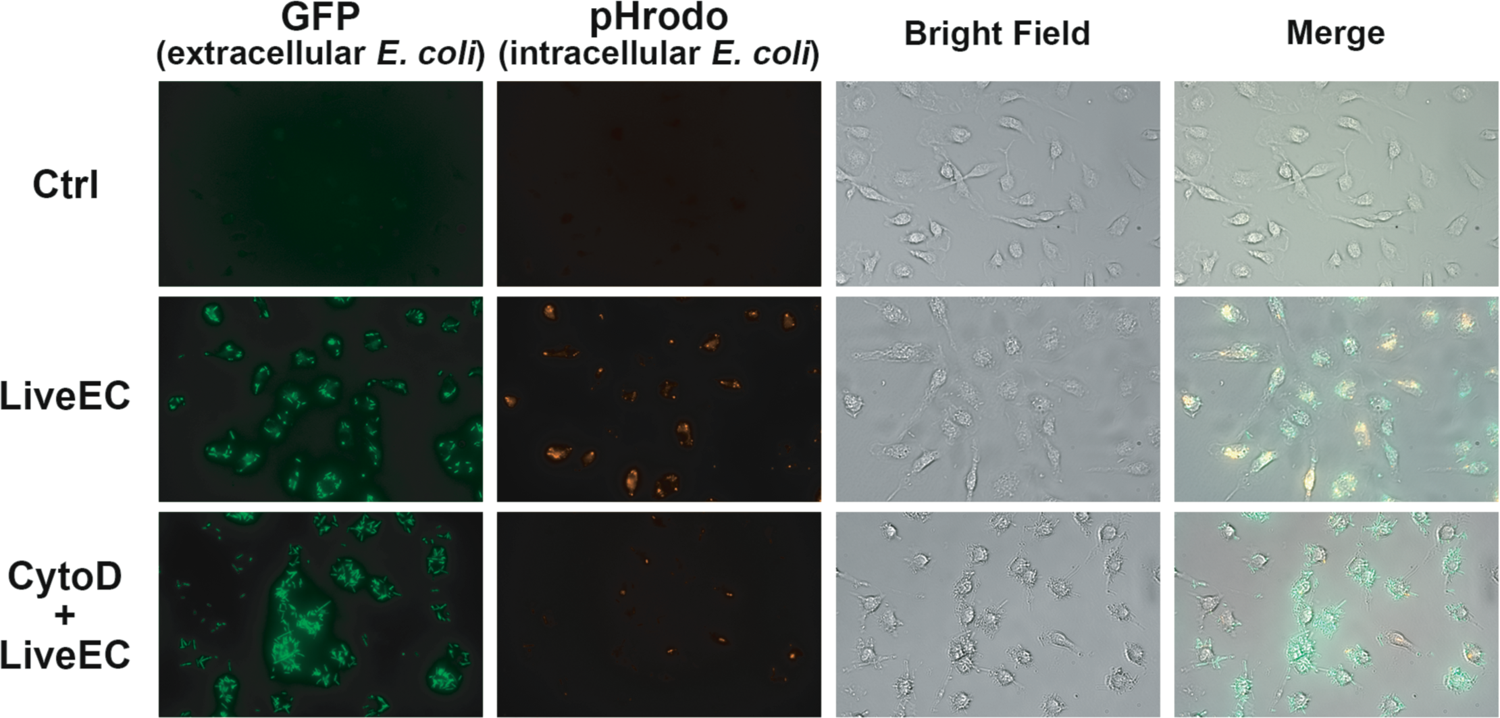
The phagocytosis inhibition efficiency in macrophages during bacterial treatment. BMDMs were incubated with pHrodo Red pre-labeled GFP-expressing *E. coli* BL21 (LiveEC) at MOI 100 in the presence or absence of phagocytosis inhibitor, Cytochalasin D (CytoD), for 30 min. Images were captured by Zeiss Axio Observer 7 40X lens after washing out the extracellular *E. coli*. Results were representative images from 3 independent experiments.

**Figure S3.**
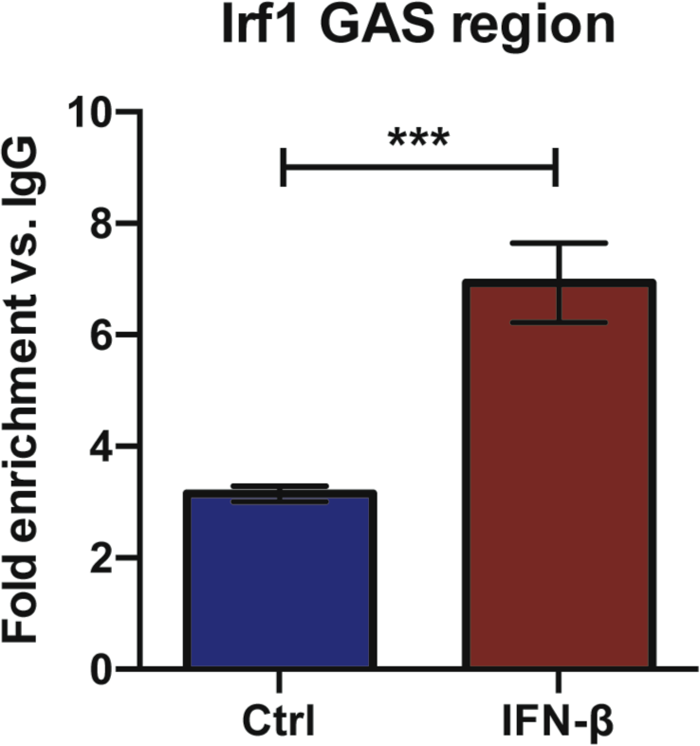
Known of STAT1 binding sites on *Irf1* GAS region in macrophages after IFN-β stimulation. *Irf1* GAS region as a positive control was used to quantify the STAT1 binding in BMDMs upon 100 U/mL IFN-β stimulation for 4 hr via qChIP assay. The results shown are combined from two independent experiments (N=2, n=4), unpaired two-tailed Student’s *t*-test was used for statistical analysis, \*\*\**p* < 0.001.

**Figure S4.**
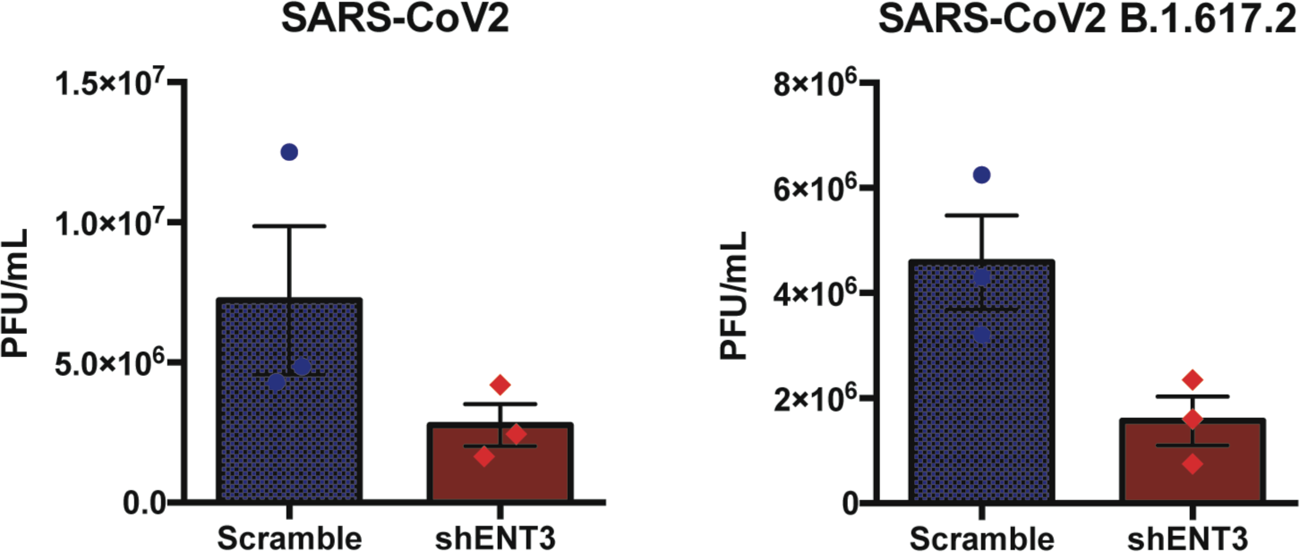
ENT3 knockdown in Calu-3 cells impaired virus production after SARS-CoV-2 infection. ENT3 knockdown Calu-3 cells were infected with SARS-CoV-2 original strain **(A)** or delta variant **(B)** at MOI=0.1 for 1 hr, and subjected for the viral titer analysis at 24 hr post-infection by plaque assay. Shown are combined results from three independent experiments (N=3).

## References

1. Aherne CM, Collins CB, Rapp CR, Olli KE, Perrenoud L, Jedlicka P, Bowser JL, Mills TW, Karmouty-Quintana H, Blackburn MR & Eltzschig HK (2018) Coordination of ENT2-dependent adenosine transport and signaling dampens mucosal inflammation. JCI Insight 3:

2. Anzilotti C, Swan DJ, Boisson B, Deobagkar-Lele M, Oliveira C, Chabosseau P, Engelhardt KR, Xu X, Chen R, Alvarez L, Berlinguer-Palmini R, Bull KR, Cawthorne E, Cribbs AP, Crockford TL, Dang TS, Fearn A, Fenech EJ, Jong SJ, Lagerholm BC, et al (2019) An essential role for the Zn2+ transporter ZIP7 in B cell development. Nature Immunology: 1–20

3. Baldwin SA (2005) Functional Characterization of Novel Human and Mouse Equilibrative Nucleoside Transporters (hENT3 and mENT3) Located in Intracellular Membranes. Journal of Biological Chemistry 280: 15880–15887

4. Bolze A, Abhyankar A, Grant AV, Patel B, Yadav R, Byun M, Caillez D, Emile J-F, Pastor-Anglada M, Abel L, Puel A, Govindarajan R, de Pontual L & Casanova J-L (2012) A mild form of SLC29A3 disorder: a frameshift deletion leads to the paradoxical translation of an otherwise noncoding mRNA splice variant. PLoS ONE 7: e29708

5. Brandenburg B, Lee LY, Lakadamyali M, Rust MJ, Zhuang X & Hogle JM (2007) Imaging poliovirus entry in live cells. PLoS Biol 5: e183

6. Campeau PM, Lu JT, Sule G, Jiang M-M, Bae Y, Madan S, Högler W, Shaw NJ, Mumm S, Gibbs RA, Whyte MP & Lee BH (2012) Whole-exome sequencing identifies mutations in the nucleoside transporter gene SLC29A3 in dysosteosclerosis, a form of osteopetrosis. Human Molecular Genetics 21: 4904–4909

7. Chakraborty M, Chu K, Shrestha A, Revelo XS, Zhang X, Gold MJ, Khan S, Lee M, Huang C, Akbari M, Barrow F, Chan YT, Lei H, Kotoulas NK, Jovel J, Pastrello C, Kotlyar M, Goh C, Michelakis E, Clemente-Casares X, et al (2021) Mechanical Stiffness Controls Dendritic Cell Metabolism and Function. CellReports 34: 108609

8. Chang C-H & Pearce EL (2016) Emerging concepts of T cell metabolism as a target of immunotherapy. Nature Immunology 17: 364–368

9. Chang C-H, Qiu J, O’Sullivan D, Buck MD, Noguchi T, Curtis JD, Chen Q, Gindin M, Gubin MM, van der Windt GJW, Tonc E, Schreiber RD, Pearce EJ & Pearce EL (2015) Metabolic Competition in the Tumor Microenvironment Is a Driver of Cancer Progression. Cell 162: 1229–1241

10. Chen Y-L, Chen T-T, Pai L-M, Wesoly J, Bluyssen HAR & Lee C-K (2013) A type I IFN-Flt3 ligand axis augments plasmacytoid dendritic cell development from common lymphoid progenitors. Journal of Experimental Medicine 210: 2515– 2522

11. Cheng Y-W, Chao T-L, Li C-L, Wang S-H, Kao H-C, Tsai Y-M, Wang H-Y, Hsieh C-L, Lin Y-Y, Chen P-J, Chang S-Y & Yeh S-H (2021) D614G Substitution of SARS-CoV-2 Spike Protein Increases Syncytium Formation and Virus Titer via Enhanced Furin-Mediated Spike Cleavage. mBio 12: e0058721

12. Cliffe ST, Kramer JM, Hussain K, Robben JH, de Jong EK, de Brouwer AP, Nibbeling E, Kamsteeg EJ, Wong M, Prendiville J, James C, Padidela R, Becknell C, van Bokhoven H, Deen PMT, Hennekam RCM, Lindeman R, Schenck A, Roscioli T & Buckley MF (2009) SLC29A3 gene is mutated in pigmented hypertrichosis with insulin-dependent diabetes mellitus syndrome and interacts with the insulin signaling pathway. Human Molecular Genetics 18: 2257–2265

13. Dufurrena Q, Amjad FM, Scherer PE, Weiss LM, Nagajyothi J, Roth J, Tanowitz HB & Kuliawat R (2017) Alterations in pancreatic β cell function and Trypanosoma cruzi infection: evidence from human and animal studies.: 1– 12

14. Durbin JE, Hackenmiller R, Simon MC & Levy DE (1996) Targeted disruption of the mouse Stat1 gene results in compromised innate immunity to viral disease. Cell 84: 443–450

15. Elbarbary NS, Tjora E, Molnes J, Lie BA, Habib MA, Salem MA & Njølstad PR (2012) An Egyptian family with H syndrome due to a novel mutation in SLC29A3 illustrating overlapping features with pigmented hypertrichotic dermatosis with insulin**-**dependent diabetes and Faisalabad histiocytosis. Pediatric Diabetes 14: 466–472

16. Freemerman AJ, Johnson AR, Sacks GN, Milner JJ, Kirk EL, Troester MA, Macintyre AN, Goraksha-Hicks P, Rathmell JC & Makowski L (2014) Metabolic Reprogramming of Macrophages: GLUCOSE TRANSPORTER 1 (GLUT1)-MEDIATED GLUCOSE METABOLISM DRIVES A PROINFLAMMATORY PHENOTYPE. J. Biol. Chem. 289: 7884–7896

17. Hall SC, Smith DR, Dyavar SR, Wyatt TA, Samuelson DR, Bailey KL & Knoell DL (2021) Critical Role of Zinc Transporter (ZIP8) in Myeloid Innate Immune Cell Function and the Host Response against Bacterial Pneumonia. J. Immunol. 207: 1357–1370

18. Hou B, Reizis B & DeFranco AL (2008) Toll-like Receptors Activate Innate and Adaptive Immunity by using Dendritic Cell-Intrinsic and -Extrinsic Mechanisms. Immunity 29: 272–282

19. Hover S, Foster B, Fontana J, Kohl A, Goldstein SAN, Barr JN & Mankouri J (2018) Bunyavirus requirement for endosomal K+ reveals new roles of cellular ion channels during infection. PLoS Pathog 14: e1006845

20. Howie D, Bokum Ten A, Necula AS, Cobbold SP & Waldmann H (2018) The Role of Lipid Metabolism in T Lymphocyte Differentiation and Survival. Front. Immunol. 8: 451–12

21. Hsu CL & Dzhagalov IL (2019) Metabolite transporters—the gatekeepers for T cell metabolism. immunometabolism: e190012

22. Hsu CL, Lin W, Seshasayee D, Chen YH, Ding X, Lin Z, Suto E, Huang Z, Lee WP, Park H, Xu M, Sun M, Rangell L, Lutman JL, Ulufatu S, Stefanich E, Chalouni C, Sagolla M, Diehl L, Fielder P, et al (2012c) Equilibrative Nucleoside Transporter 3 Deficiency Perturbs Lysosome Function and Macrophage Homeostasis. Science 335: 89–92

23. Jaouadi H, Zaouak A, Sellami K, Messaoud O, Chargui M, Hammami H, Jones M, Jouini R, Chadli Debbiche A, Chraiet K, Fenniche S, Mrad R, Mokni M, Turki H, Benkhalifa R & Abdelhak S (2018) H syndrome: Clinical, histological and genetic investigation in Tunisian patients. J Dermatol 45: 978–985

24. Jordheim LP, Durantel D, Zoulim F & Dumontet C (2013) Advances in the development of nucleoside and nucleotide analogues for cancer and viral diseases. Nature Publishing Group 12: 447–464

25. Kelly B & O’Neill LA (2015) Metabolic reprogramming in macrophages and dendritic cells in innate immunity. Nature Publishing Group 25: 771–784

26. Landfear SM (2011) Nutrient Transport and Pathogenesis in Selected Parasitic Protozoa. Eukaryotic Cell 10: 483–493

27. Lu Y-C, Yeh W-C & Ohashi PS (2008) LPS/TLR4 signal transduction pathway. Cytokine 42: 145–151

28. Macanas-Pirard P, Broekhuizen R, González A, Oyanadel C, Ernst D, García P, Montecinos VP, Court F, Ocqueteau M, Ramirez P & Nervi B (2017) Resistance of leukemia cells to cytarabine chemotherapy is mediated by bone marrow stroma, involves cell-surface equilibrative nucleoside transporter-1 removal and correlates with patient outcome. Oncotarget 8: 23073–23086

29. Moon J-S, Hisata S, Park M-A, DeNicola GM, Ryter SW, Nakahira K & Choi AMK (2015) mTORC1-Induced HK1-Dependent Glycolysis Regulates NLRP3 Inflammasome Activation. CellReports 12: 102–115

30. Morgan NV, Morris MR, Cangul H, Gleeson D, Straatman-Iwanowska A, Davies N, Keenan S, Pasha S, Rahman F, Gentle D, Vreeswijk MPG, Devilee P, Knowles MA, Ceylaner S, Trembath RC, Dalence C, Kismet E, Köseoğlu V, Rossbach H-C, Gissen P, et al (2010) Mutations in SLC29A3, encoding an equilibrative nucleoside transporter ENT3, cause a familial histiocytosis syndrome (Faisalabad histiocytosis) and familial Rosai-Dorfman disease. PLoS Genet 6: e1000833

31. Mudhakir D & Harashima H (2009) Learning from the viral journey: how to enter cells and how to overcome intracellular barriers to reach the nucleus. AAPS J 11: 65–77

32. Nair S, Strohecker AM, Persaud AK, Bissa B, Muruganandan S, McElroy C, Pathak R, Williams M, Raj R, Kaddoumi A, Sparreboom A, Beedle AM & Govindarajan R (2019) Adult stem cell deficits drive Slc29a3 disorders in mice. Nature Communications: 1–20

33. O’Neill LAJ & Pearce EJ (2015) Immunometabolism governs dendritic cell and macrophage function. J. Exp. Med. 64: jem.20151570–9

34. Platanias LC (2005) Mechanisms of type-I- and type-II-interferon-mediated signalling. Nature Reviews Immunology 5: 375–386

35. Priya TP, Philip N, Molho-Pessach V, Busa T, Dalal A & Zlotogorski A (2010) H syndrome: novel and recurrent mutations in SLC29A3. British Journal of Dermatology 162: 1132–1134

36. Ryu W-S (2017) Virus life cycle. Molecular Virology of Human Pathogenic Viruses: 31–45

37. Schneider WM, Chevillotte MD & Rice CM (2014) Interferon-stimulated genes: a complex web of host defenses. Annu. Rev. Immunol. 32: 513–545

38. Shibata T, Taoka M, Saitoh SI, Yamauchi Y, bioRxiv YM 2019 Nucleosides drive histiocytosis in SLC29A3 disorders by activating TLR7*. biorxiv.org*

39. Suganuma T & Workman JL (2021) Nucleotide Metabolism Behind Epigenetics. Front. Endocrinol. 12: 731648

40. Takagaki K, Katsuma S, Kaminishi Y, Horio T, Nakagawa S, Tanaka T, Ohgi T & Yano J (2004) Gene-expression profiling reveals down-regulation of equilibrative nucleoside transporter 1 (ENT1) in Ara-C-resistant CCRF-CEM-derived cells. Journal of Biochemistry 136: 733–740

41. Tang T, Li L, Tang J, Li Y, Lin WY, Martin F, Grant D, Solloway M, Parker L, Ye W, Forrest W, Ghilardi N, Oravecz T, Platt KA, Rice DS, Hansen GM, Abuin A, Eberhart DE, Godowski P, Holt KH, et al (2010) A mouse knockout library for secreted and transmembrane proteins. Nature Biotechnology 28: 749– 755

42. Vitale I, Manic G, Coussens LM, Kroemer G & Galluzzi L (2019) Macrophages and Metabolism in the Tumor Microenvironment. Cell Metabolism 30: 36–50

43. Wang W & Zou W (2020) Amino Acids and Their Transporters in T Cell Immunity and Cancer Therapy. Molecular Cell 80: 384–395

44. Wang W, Xu L, Su J, Peppelenbosch MP & Pan Q (2017) Transcriptional Regulation of Antiviral Interferon-Stimulated Genes. Trends in Microbiology 25: 573–584

45. Wei C-W, Lee C-Y, Lee D-J, Chu C-F, Wang J-C, Wang T-C, Jane W-N, Chang Z-F, Leu C-M, Dzhagalov IL & Hsu C-L (2018) Equilibrative Nucleoside Transporter 3 Regulates T Cell Homeostasis by Coordinating Lysosomal Function with Nucleoside Availability. CellReports 23: 2330–2341

46. Winstel V, Missiakas D & Schneewind O (2018) Staphylococcus aureus targets the purine salvage pathway to kill phagocytes. Proc. Natl. Acad. Sci. U.S.A. 115: 6846–6851

47. Wu K-C, Lee C-Y, Chou F-Y, Chern Y & Lin C-J (2020) Deletion of equilibrative nucleoside transporter-2 protects against lipopolysaccharide-induced neuroinflammation and blood-brain barrier dysfunction in mice. Brain Behavior and Immunity 84: 59–71

48. Young JD, Yao SYM, Baldwin JM, Cass CE & Baldwin SA (2013a) The human concentrative and equilibrative nucleoside transporter families, SLC28 and SLC29. Mol. Aspects Med. 34: 529–547

49. Young JD, Yao SYM, Baldwin JM, Cass CE & Baldwin SA (2013b) Molecular Aspects of Medicine. Molecular Aspects of Medicine 34: 529–547

